# Mast cell-derived BH4 is a critical mediator of postoperative pain

**DOI:** 10.1101/2023.01.24.525378

**Authors:** Philipp Starkl, Gustav Jonsson, Tyler Artner, Bruna Lenfers Turnes, Nadine Serhan, Tiago Oliveira, Laura-Marie Gail, Karel Stejskal, Keith M. Channon, Thomas Köcher, Georg Stary, Victoria Klang, Nicolas Gaudenzio, Sylvia Knapp, Clifford J. Woolf, Josef M. Penninger, Shane J.F. Cronin

## Abstract

Postoperative pain affects most patients after major surgery and can transition to chronic pain. Here, we discovered that postoperative pain hypersensitivity correlated with markedly increased local levels of the metabolite BH4. Gene transcription and reporter mouse analyses after skin injury identified neutrophils, macrophages and mast cells as primary postoperative sources of GTP cyclohydrolase-1 (*Gch1*) expression, the rate-limiting enzyme in BH4 production. While specific *Gch1* deficiency in neutrophils or macrophages had no effect, mice deficient in mast cells or mast cell-specific *Gch1* showed drastically decreased postoperative pain after surgery. Skin injury induced the nociceptive neuropeptide substance P, which directly triggers the release of BH4-dependent serotonin in mouse and human mast cells. Substance P receptor blockade substantially ameliorated postoperative pain. Our findings underline the unique position of mast cells at the neuro-immune interface and highlight substance P-driven mast cell BH4 production as promising therapeutic targets for the treatment of postoperative pain.

## Introduction

Each year a quarter of a billion people undergo surgery, with the vast majority experiencing acute postoperative pain (Gan, 2017; Weiser et al., 2008). Postoperative pain causes suffering, delays recovery, and can transition into chronic pain, causing a massive economic burden on individuals and the healthcare sector (Gan, 2017; Kehlet et al., 2006; Weiser et al., 2008). The intensity of acute pain is a major predictor for the chronification of postoperative pain (Fletcher et al., 2015). Current treatments of acute postoperative pain, such as opioids, ketamine or non-steroidal anti-inflammatory drugs (NSAIDs) have undesirable side effects, can be ineffective and bear the risk of enormous additional socioeconomic and healthcare burden upon misuse (Dolin et al., 2002; Foley, 2006; Shipton et al., 2018). Thorough characterization of the molecular and cellular mechanisms involved in postoperative pain hypersensitivity could facilitate the development of novel therapeutic strategies.

Tetrahydrobiopterin, BH4, is an essential cofactor for several enzymes with critical physiologic and metabolic functions, including nitric oxide (NO) production and synthesis of amine neurotransmitters, such as norepinephrine, epinephrine, serotonin and dopamine (Werner et al., 2011). BH4 production is controlled by GTP cyclohydrolase-1 (hereafter referred to as GCH1), which is the first and rate-limiting enzyme of the *de novo* BH4 biosynthesis pathway. We and others have demonstrated a close correlation between GCH1 and BH4 levels in injured nerves and pain intensity in animal models and in humans suffering from chronic pain (Belfer et al., 2015; Cronin et al., 2022; Fujita et al., 2020; Latremoliere et al., 2015; Nasser and Moller, 2014; Tegeder et al., 2006). Furthermore, BH4 has important functions in various immune cells, including macrophages (McNeill et al., 2018), neutrophils (Nagarkoti et al., 2019), and T cells (Cronin et al., 2018).

Surgical tissue injury causes inflammation, characterized by the infiltration and activation of immune cells (Arias et al., 2009). Inflammatory mediators produced during this process, such as interleukin (IL)-1ß, IL-6 or tumor necrosis factor-alpha (TNFα) cause pain-like behaviors by direct activation of pain-initiating sensory nerves (nociceptors) (Baral et al., 2019). On the other hand, activated nociceptors release chemotactic compounds that recruit and/or activate immune cells (Tauber et al., 2021), establishing a potential reciprocal interaction between the nervous and immune systems in postoperative pain. Improved knowledge of such neuro-immune crosstalk is required to fully understand the pathophysiology associated with postsurgical pain hypersensitivity. Here, we chart this crosstalk and identify an immune cell-specific GCH1/BH4 pathway in mast cells as a critical regulator of pain hypersensitivity after surgical injury.

## Results

### Skin immune cells express *Gch1* after surgical tissue damage

To explore postoperative pain, we performed an incision of the hind paw skin in mice with a tearing of the underlying skeletal muscle, as described previously (Brennan et al., 1996; Pogatzki and Raja, 2003). The subsequent pain hypersensitivity is a result of the injury-induced inflammation (Ghasemlou et al., 2015) as well as nerve damage (Hill et al., 2010). We observed that incision-mediated mechanical pain sensitivity peaked after 24 hours and returned to baseline threshold levels after ∼96 hours (**Figure 1A and B**). Similarly, we observed, thermal hyperalgesia at 24 hours after incision, which returned to baseline after 48 hours. Using the 24-hour time point after incision, when pain hypersensitivity was strongest, we observed a significant increase in paw thickness in the incised (ipsilateral) compared to the control (contralateral) paw (**Figure 1C**). GCH1 is the rate-limiting enzyme in *de novo* BH4 biosynthesis and contributes to neuropathic pain by driving increased BH4 levels in nerves and dorsal root ganglia (DRG) after nerve injury as observed in the spared nerve injury (SNI) neuropathic pain rodent model (Decosterd and Woolf, 2000; Fujita et al., 2020; Latremoliere et al., 2015; Tegeder et al., 2006). Incision injury increased BH4 levels in the incised paw but not in the respective sciatic nerve in wild type mice (**Figure 1D, E**). Whereas SNI enhanced sciatic nerve BH4 levels (**Figure 1E**) and *Gch1* expression in the ipsilateral DRGs, such *Gch1* induction was not observed following surgical injury, as determined by *Gch1*-GFP reporter mice (**Figure 1F and Figure S1A**) (Latremoliere et al., 2015).

**Figure 1.**
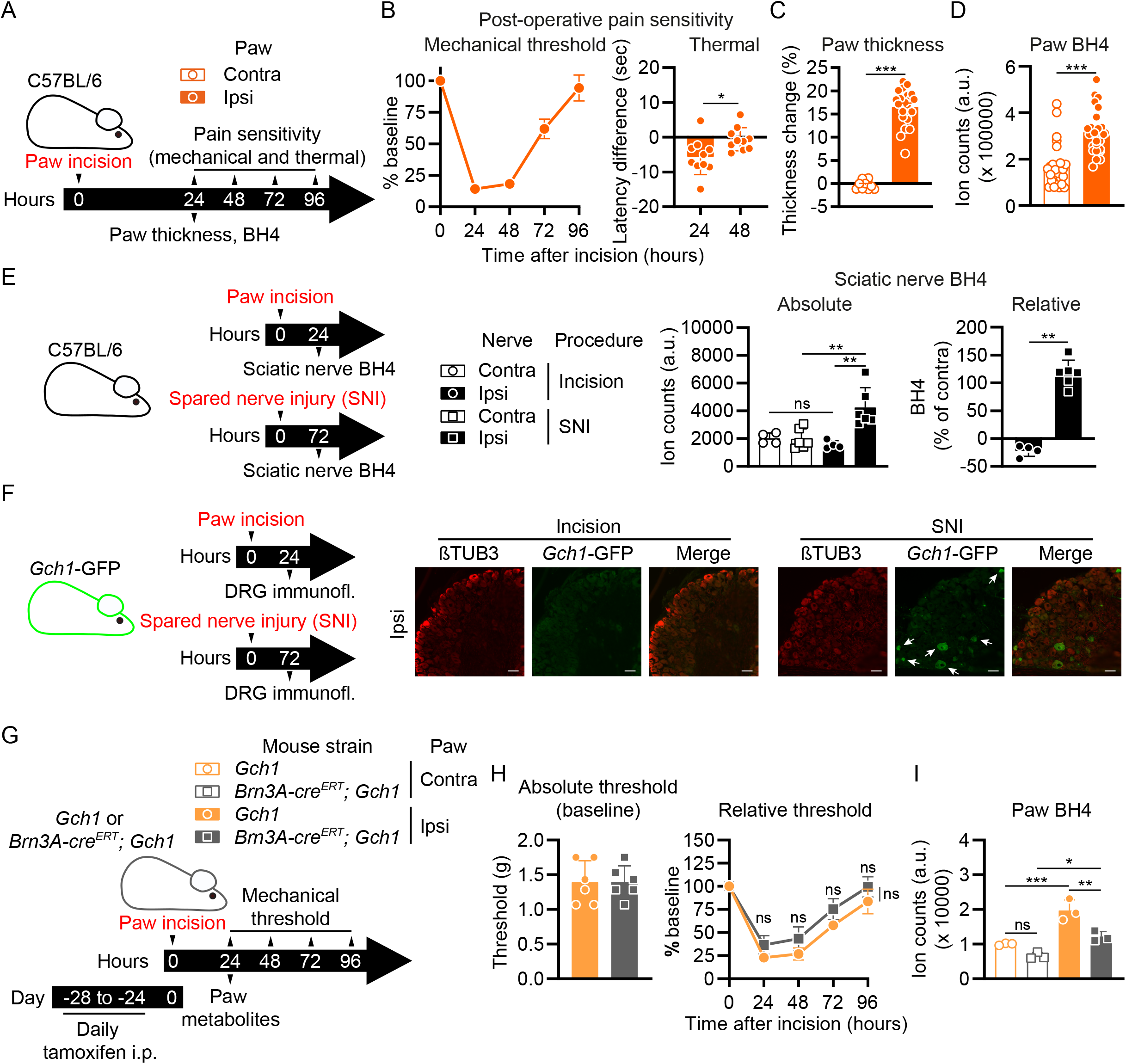
Tissue damage and pain-like hypersensitivity is associated with *Gch1*-expressing immune cells in the incision-injured paw. (A) Experimental scheme for (B to D). C57BL/6 wild type mice were subjected to paw incision. Contralateral (Contra) and ipsilateral (Ipsi) paw mechanical pain sensitivity and paw BH4 levels were assessed at the indicated time points after incision. (B) Time course of mechanical pain sensitivity relative to baseline at the indicated time points after incision (left panel; n=28) and thermal pain sensitivity 24 and 48 hours after incision compared to baseline before incision (right panel; n=10). (C) Thickness change (compared to baseline) of Contra and Ipsi paws 24 hours after incision (n=10-21). (D) BH4 levels in Contra and Ipsi paws 24 hours after incision (n=26). (E) Experimental scheme and measurement of BH4 levels in the Contra and Ipsi L3-L4 dorsal root ganglion (DRG) tissues of incision injury- (n=4) wild type mice and a spared nerve injury (SNI)-treated (n=6) wild type mouse as control. (F) Experimental scheme and representative immunofluorescence (immunofl.) images of anti-beta-TUBULIN-III (ßTUB3) and anti-GFP (*Gch1*-GFP) staining in the Ipsi L3-L4 DRG tissues of incision injury- (representative of n=3) and spared nerve injury (SNI)-treated (representative of n=3) *Gch1*-GFP reporter mice. Arrows indicate *Gch1*-GFP-positive cells. Scale bars represent 50 μm. (G) Experimental scheme for (H and I). *Gch1*^*flox/flox*^ (*Gch1*) control mice or *Gch1* mice expressing tamoxifen-inducible cre recombinase in sensory neurons (*Brn3A-cre*^*ERT*^; *Gch1*^*flox/flox*^; *Brn3A-cre*^*ERT*^; *Gch1*) mice were treated by daily intraperitoneal tamoxifen injections (2mg/mouse) 28 – 24 days before paw incision. Contra and Ipsi paw mechanical pain sensitivity and paw BH4 levels were assessed at the indicated time points after incision. (H) Absolute mechanical threshold at baseline and relative (to baseline) mechanical threshold kinetics (n=6). (I) Paw BH4 levels (n=3). (B to D, H) Mann-Whitney test; (E, I) One-Way ANOVA with Tukey’s multiple comparisons test; (H) Two-Way ANOVA with Sidak’s multiple comparisons test (comparing individual time points) and Two-Way ANOVA with repeated measures with Geisser-Greenhouse correction (overall comparison); * P ≤ 0.05, ** P ≤ 0.01, *** P ≤ 0.001, ns – not significant. Error bars indicate (B to E) SEM or (H and I) SD. Symbols in bar graphs represent individual mice.

To determine whether sensory neuron-derived BH4 contributes to the pain hypersensitivity after paw incision, we deleted *Gch1* in DRG neurons in adult mice using a tamoxifen inducible Cre line, *Brn3A-Cre*^*ERT*^ (Shacham-Silverberg et al., 2018) combined with floxed *Gch1* alleles (*Gch1*^*flox/flox*^ mice, hereafter referred to as *Gch1*) (Chuaiphichai et al., 2014; Latremoliere et al., 2015) (**Figure 1G**). There were no significant differences in mechanical hypersensitivity between control and *Brn3A-cre*^*ERT*^; *Gch1* animals after incision (**Figure 1H**). Interestingly, mice with *Gch1*-deficient sensory neurons showed substantially less BH4 in the incised paw compared to control animals (**Figure 1I**), indicating that neuronal BH4 contributes to the local increase of the metabolite after injury but not to the pain hypersensitivity. Altogether, these data show that paw surgery leads to locally-increased BH4 which correlates with enhanced pain sensitivity. However, while sensory neurons contribute to elevated BH4, *Gch1* deficiency in these neurons apparently did not influence mechanical allodynia after surgical incision.

Besides its role in neuronal physiology (Werner et al., 2011), BH4 exerts important immune functions, including NO production in macrophages (McNeill et al., 2018) and cofactor-independent roles in T cell proliferation by regulating mitochondrial bioenergetics (Cronin et al., 2018). To assess whether immune cells could be an additional source of BH4 after surgery and contribute to postoperative pain, we screened *Gch1* expression in various immune cell populations using publicly available gene expression data (Heng et al., 2008; Seita et al., 2012; Wu et al., 2009). We found that *Gch1* is expressed in a plethora of immune cell types, including dendritic cells (DCs), natural killer (NK) cells, neutrophils, type 1 innate lymphoid cells (ILC1s), ILC3s, eosinophils, macrophages and mast cells (**Figure 2A; Figure S1B and Table S1**). Using these data sets as a guide, we next aimed to identify *Gch1*-expressing immune cells after surgery and found increased leukocyte *Gch1* expression in the incised paw of *Gch1*-GFP reporter mice (**Figure 2B; Figure S1C and S1D)**. Among the immune cell populations meeting two thresholds (threshold 1: more than 1 cell with *Gch1*-GFP signal above autofluorescence of respective wild type controls; threshold 2: representing more than 1% of all *Gch1*-GFP^+^ leukocytes; **Figure S1D**), we prioritized neutrophils, macrophages and mast cells as potential candidates for surgery-induced cellular BH4 production (**Figure 2C**).

**Figure 2.**
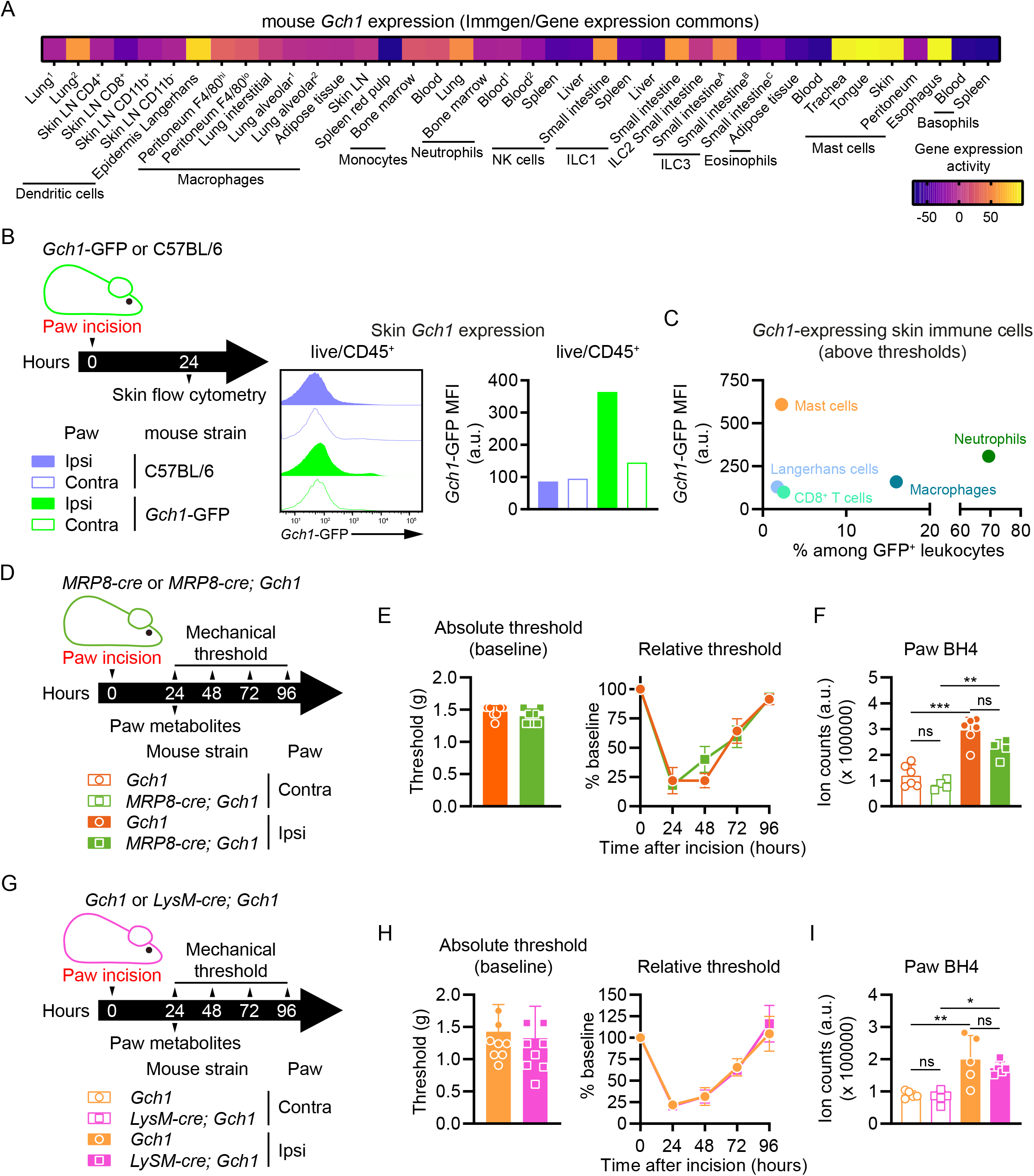
Myeloid immune cells express *Gch1* in injured skin. (A) Mouse *Gch1* gene expression levels in immune cell populations (n=1-4) derived from the *ImmGen* Microarray dataset using the *Gene Expression Commons* platform (Table S1). (B and C) Twenty-four hours after incision, skin cells from Contra and Ipsi paws of *Gch1*-GFP reporter (one pool of 5) or wild type C57BL/6 (one pool of 5) mice were analyzed by flow cytometry. (B) Left panel: experimental scheme; middle panel: histograms depicting *Gch1*-GFP signal of live CD45^+^ cells; right panel: *Gch1*-GFP mean fluorescence intensity (MFI) signals of live CD45^+^ cells. (C) Representation of immune cell populations meeting threshold criteria (see methods and Fig. S1), depicting *Gch1*-GFP MFI (y-axis) and % among GFP^+^ leukocytes (x-axis). (D) Experimental scheme for (E and F). Control *Gch1*^*flox/flox*^ (*Gch1*) mice and animals with neutrophil-specific *Gch1* deficiency (*MRP8-cre; Gch1)* were subjected to incision injury. Contralateral (Contra) and ipsilateral (Ipsi) paw mechanical pain sensitivity and BH4 levels were assessed 24 hours after incision. (E) Absolute mechanical threshold at baseline and relative (to baseline) mechanical threshold kinetics (n=7). (F) Paw BH4 levels (n=4-6). (G) Experimental scheme for (H and I). *Gch1*^*flox/flox*^ (*Gch1*) mice or *Gch1* mice expressing cre recombinase under control of the lysozyme promoter in macrophages (as well as in proportions of neutrophils and monocytes; *LyzM-cre; Gch1*) mice were treated by paw incision. Contra and Ipsi paw mechanical pain sensitivity and paw BH4 levels were assessed at indicated time points after incision. (H) Absolute mechanical threshold at baseline and relative (to baseline) mechanical threshold kinetics (n=10). (I) Paw BH4 levels (n=5). (F, I) One-Way ANOVA with Tukey’s multiple comparisons test; * P ≤ 0.05, ** P ≤ 0.01, *** P ≤ 0.001, ns – not significant. Error bars indicate SD. Symbols in bar graphs represent individual mice.

### Mast cells contribute to postoperative pain hypersensitivity

Neutrophils represented the subtype with the highest proportion (69.5%) among *Gch1*-expressing leukocytes in the incised paw (**Figure 2C; Figure S1D**). To investigate their role in incision-induced pain, we generated *MRP8-cre; DTR* mice in which the diphtheria toxin receptor (DTR) is specifically expressed by neutrophils (Reber et al., 2017). Administration of diphtheria toxin (DT) resulted in efficient neutrophil depletion (**Figure S2A**). However, we observed no differences in mechanical allodynia between DT-treated control and *MRP8-cre; DTR* mice (**Figure S2B and S2C**). Moreover, a neutrophil deficiency did not affect paw BH4 levels (**Figure S2D**). We also used mice with a neutrophil-specific *Gch1* knockout (*MRP8-cre; Gch1*) to directly assess if there is any contribution of neutrophil-derived GCH1/BH4 to postoperative pain (**Figure 2D**). Animals with *Gch1*-deficient neutrophils and their respective controls displayed comparable mechanical pain hypersensitivity as well as paw BH4 levels before and after injury (**Figure 2E and 2F**). These data are consistent with previous reports of only minor neutrophil contributions to incision-induced pain hypersensitivity (Ghasemlou et al., 2015; Segelcke et al., 2021). To analyze the potential contribution of macrophage-derived BH4 to postoperative pain, we investigated mechanical hypersensitivity and paw metabolites in *LysM-cre*; *Gch1* mice (with a Cre-mediated *Gch1* deletion in macrophages as well as neutrophils and monocytes (Abram et al., 2014; Imai et al., 2008)) (**Figure 2G**). Ablation of *Gch1* in *LysM*-expressing cells did not influence pain hypersensitivity (**Figure 2H**) nor paw BH4 (**Figure 2I**) after incision.

Although neutrophils and macrophages constitute the cell types with the highest proportion of *Gch1*^+^ leukocytes, mast cells exhibited the highest *Gch1* expression (**Figure 2C; FigureS1D**). In addition, we observed *Gch1*-expressing mast cells in the injured paw of *Gch1*-GFP reporter mice (**Figure 3A**). Upon appropriate activation, mast cells rapidly release cytoplasmic granules loaded with various preformed bioactive compounds (Mukai et al., 2018; St John et al., 2022). Two of the major mediators stored in mast cell granules are serotonin and histamine, which induce pain and itch (Rosa and Fantozzi, 2013; Sommer, 2004). Serotonin is of particular interest since BH4 is a necessary cofactor for serotonin synthesis from tryptophan by tryptophan hydroxylase (Werner et al., 2011). Interestingly, serotonin, but not histamine, levels increased significantly upon paw incision in wild type mice (**Figure 3B**), mirroring the BH4 dynamics and pain hypersensitivity. We next examined a role for mast cells in the postoperative pain hypersensitivity using mast cell-deficient *Mcpt5-cre; DTA* mice (Dudeck et al., 2011) in which expression of diphtheria toxin subunit A (*DTA*) is driven by the connective tissue mast cell-specific promoter *mast cell protease 5* (*Mcpt5*) (**Figure 3C; Figure S3A**). Strikingly, mast cell-deficient *Mcpt5-cre; DTA* mice exhibited decreased pain hypersensitivity (**Figure 3D**). This observation was accompanied by a marked reduction in serotonin and histamine levels (**Figure 3E**) while paw thickness was not affected (**Figure S3B**). These results demonstrate a prominent role for mast cells in pain hypersensitivity after surgical tissue injury.

**Figure 3.**
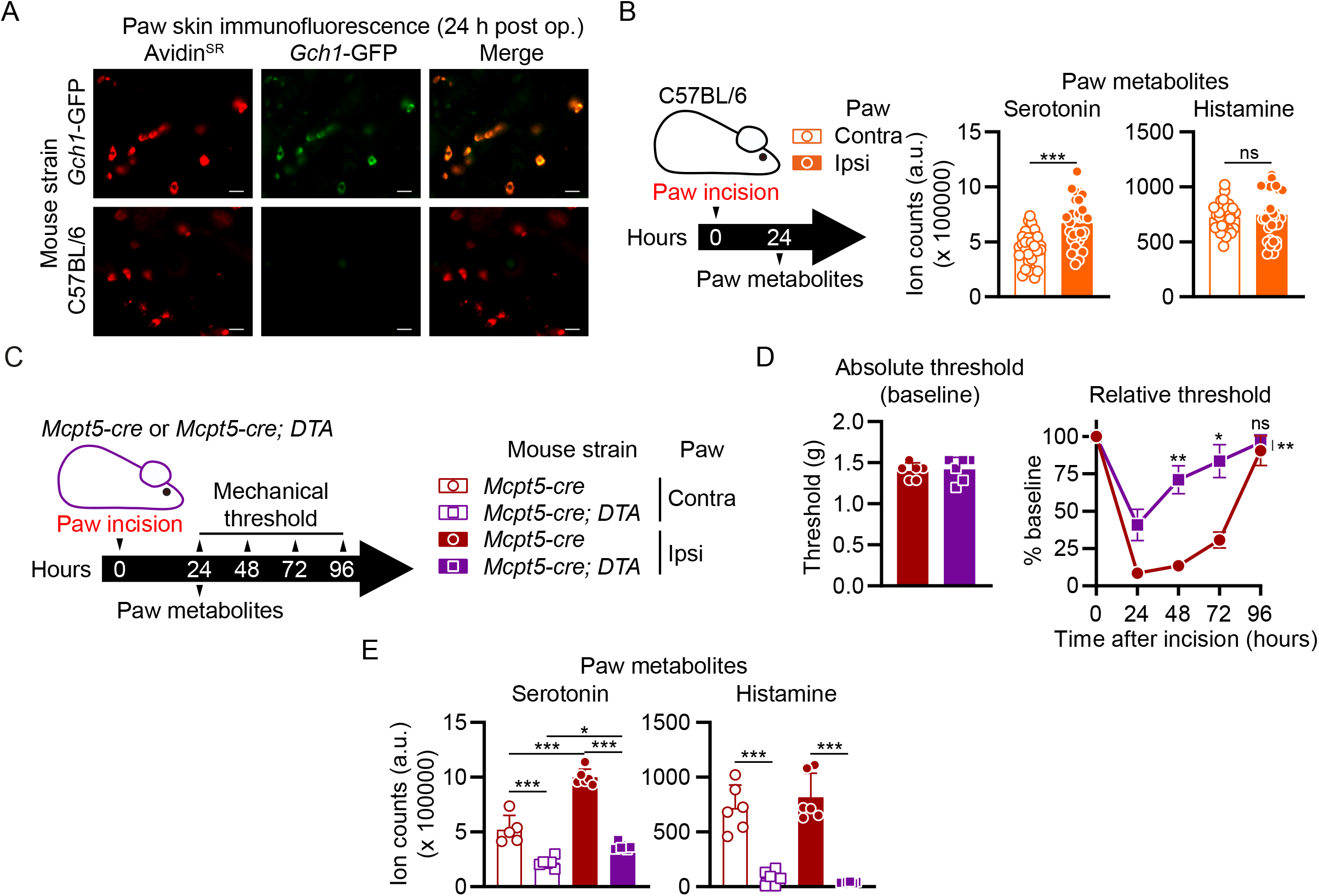
Mast cells are major contributors to postoperative pain-like hypersensitivity. (A) Representative skin immunofluorescence images of sulforhodamin-101-labelled avidin (Avidin^SR^) and anti-GFP staining in Ipsi paws of wild type and *Gch1*-GFP reporter mice 24 hours post incision injury. Scale bars represent 20 µm. (B) Experimental scheme and measurements of serotonin and histamine levels in Contra and Ipsi paws 24 hours after incision injury in wild type animals (n=30-33). (C) Experimental scheme for (D and E). Postoperative pain kinetics and paw metabolites were assessed in control (*Mcpt5-cre*) and mast cell-depleted (*Mcpt5-cre; DTA*) mice. (D) Absolute mechanical threshold at baseline and relative (to baseline) mechanical threshold kinetics (n=6). (E) Paw tissue metabolites 24 hours post incision injury of Contra and Ipsi paws (n=5-6). (B) Mann-Whitney test; (E) One-Way ANOVA with Tukey’s multiple comparisons test; (D) Two-Way ANOVA with Sidak’s multiple comparisons test (comparing individual time points) and Two-Way ANOVA with repeated measures with Geisser-Greenhouse correction (overall comparison); * P ≤ 0.05, ** P ≤ 0.01, *** P ≤ 0.001, ns – not significant. Error bars indicate (B, D) SEM or (E) SD. Symbols in bar graphs represent individual mice.

### Manipulation of mast cell BH4 modulates pain hypersensitivity after surgical injury

To dissect the involvement of mast cell-derived BH4 in pain hypersensitivity, we generated *Mcpt5-cre; Gch1* mice with mast cell-specific *Gch1* deficiency (**Figure 4A**). Following the surgical injury, *Mcpt5-cre; Gch1* mice displayed markedly reduced mechanical (**Figure 4B**) as well as thermal (**Figure 4C**) hypersensitivity compared to control animals. Importantly, the reduced mechanical allodynia was observed in both male and female *Mcpt5-cre; Gch1* mice (**Figure S3C**). The reduced pain-like behaviors of *Mcpt5-cre; Gch1* mice were accompanied by lower paw BH4 and serotonin levels, whereas histamine levels remained unaffected (**Figure 4D**). Paw incision induces robust infiltration of immune cells and a significant increase in paw weight after injury (Green et al., 2019); however, mast cell-specific loss of *Gch1* did not influence paw thickness and weight (**Figure 4E and Figure S3D**) nor the numbers and distributions of infiltrating immune cells (leukocytes, neutrophils, macrophages, mast cells) after paw injury (**Figure S3E**). Having observed the drastically decreased pain hypersensitivity associated with specific mast cell BH4 deficiency, we asked whether enhancing the mast cell BH4 production machinery would increase postoperative allodynia. To test this idea, we generated mice combining the *Mcpt5-cre* line with an allele construct allowing cre-mediated *Gch1* overexpression (*Mcpt5-cre; GOE* **Figure 4F**) (Latremoliere et al., 2015). Notably, *Gch1* overexpression *per se* did not cause altered pain sensitivity (**Figure 4G**) despite substantially increased tissue BH4 and serotonin levels at baseline (**Figure 4H**). In contrast, *Mcpt5-cre; GOE* responded to paw incision with significantly greater postoperative pain hypersensitivity (**Figure 4G**) which was associated with higher amounts of skin serotonin (**Figure 4H**). Similar to mast cell-specific *Gch1* deletion, overexpression neither affected paw weight (**Figure S4A**) nor immune cell influx (**Figure S4B**).

**Figure 4.**
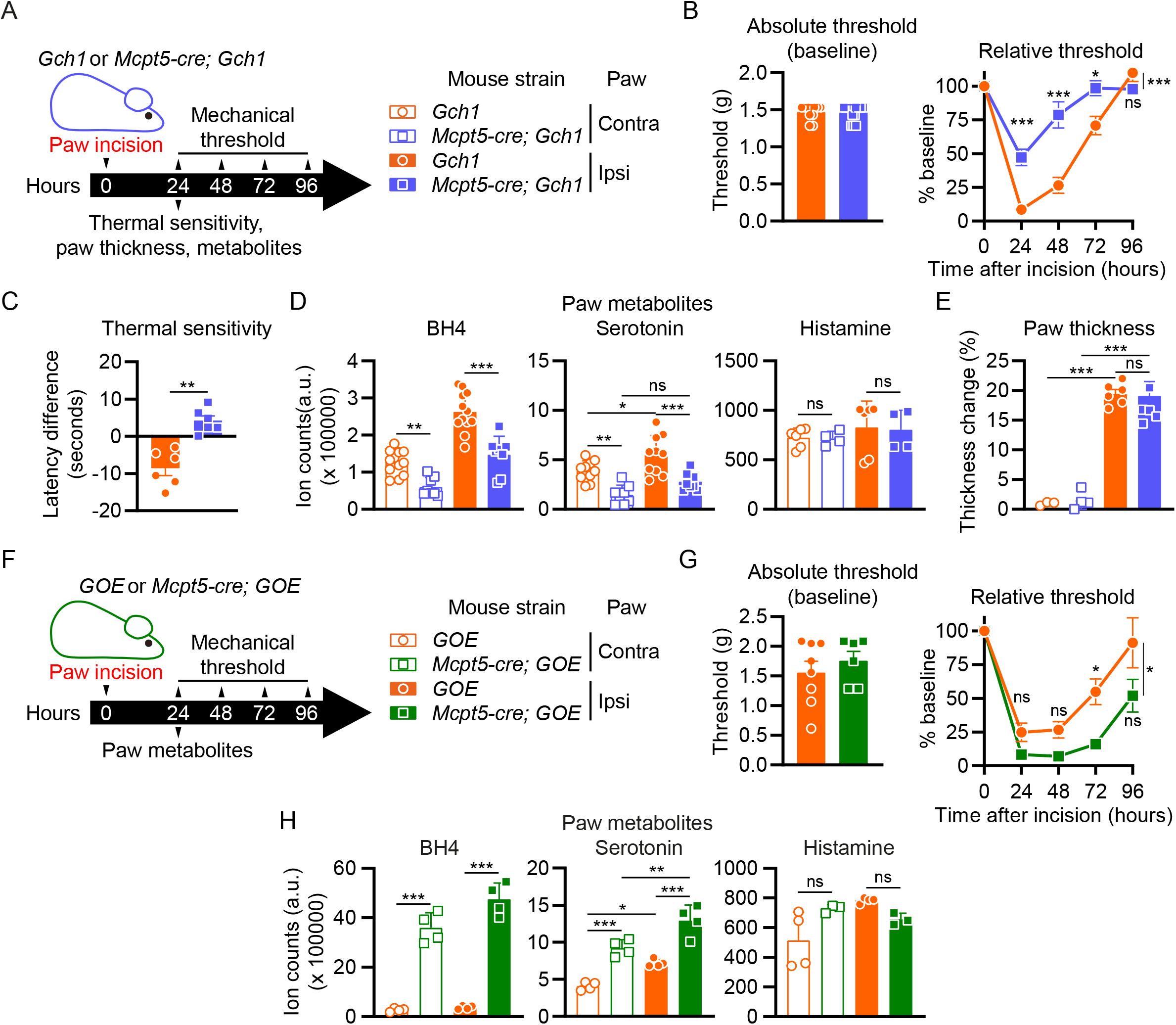
Mast cell-specific *Gch1* deletion and overexpression modulates postoperative mechanical hypersensitivity. (A) Experimental scheme for (B – E). Postoperative mechanical and thermal pain hypersensitivity as well as paw thickness and metabolites were assessed in control (*Gch1*) and *Mcpt5-cre; Gch1* mice. (B) Absolute mechanical threshold at baseline and relative (to baseline) mechanical threshold kinetics (n=11-14). (C) Thermal pain sensitivity 24 hours after incision compared to baseline before incision (n=5-6). (D) Paw tissue metabolites of Contra and Ipsi paws 24 hours post incision injury (n=4-12). (E) Contra and Ipsi paw thickness change 24 hours post incision injury relative to baseline (before incision) (n=5-7). (F) Experimental scheme for (G and H). Postoperative pain kinetics and metabolites in contralateral (Contra) and ipsilateral (Ipsi) paws in control (*GOE*) and *Mcpt5-cre; GOE* mice. (G) Absolute mechanical threshold at baseline and relative (to baseline) mechanical threshold kinetics (n=6-8). (H) Contra and Ipsi paw tissue metabolites 24 hours post incision injury (n=3-4). (B, G) Two-Way ANOVA with Sidak’s multiple comparisons test (comparing individual time points) and Two-Way ANOVA with repeated measures with Geisser-Greenhouse correction (overall comparison); (C) Mann-Whitney test; (D, E, H) One-Way ANOVA with Tukey’s multiple comparisons test; * P ≤ 0.05, ** P ≤ 0.01, *** P ≤ 0.001, ns – not significant. Error bars indicate (B, D [BH4 and serotonin panels], E, G) SEM or (C, D [histamine panel], H) SD. Symbols in bar graphs represent individual mice.

Given the critical role of mast cell BH4 in our model, we next wanted to explore the therapeutic applicability of local interference with BH4 production. We recently described a novel inhibitor of BH4 synthesis, QM385, which targets sepiapterin reductase (SPR), the terminal enzyme in the *de novo* BH4 biosynthesis pathway (Cronin et al., 2018). Intradermal QM385 administration significantly reduced skin BH4 and serotonin and elevated sepiapterin (which accumulates due to the QM385-mediated SPR blockade) after 8 hours, returning to baseline levels 24 hours after treatment (**Figure S4C**). Although a single paw injection of QM385 two hours before incision did not significantly alter mechanical allodynia, there was a trend towards an amelioration of the pain (**Figure S4D**).

Overall, our results indicate that mast cell-specific *Gch1* levels profoundly influence BH4, serotonin and mechanical allodynia upon paw incision.

### Stimulated mast cells rapidly release BH4, serotonin and their biosynthetic enzymes after activation

Nociceptors express the serotoninergic receptors 5-HT3 and 5-HT2A (Van Steenwinckel et al., 2009; Zeitz et al., 2002) and serotonin stimulation via these receptors increases neuronal excitability and pain signaling (Liu et al., 2020). Our *in vivo* and *in vitro* data indicate that BH4 levels in mast cells regulate serotonin synthesis and directly affect post-injury pain thresholds. Tamoxifen-treated *Brn3a-creERT; Gch1* mice displayed significantly reduced BH4 levels in the paw after incision but this reduction *per se* did not affect pain sensitivity (**Figure 1H**). Moreover, serotonin levels were unaffected in these animals (**Figure S5A**). To determine whether BH4 levels influence the amount of serotonin secreted by activated mast cells, we generated bone marrow-derived cultured mast cells (BMCMCs) from control, *Mcpt5-cre; Gch1* as well as *Mcpt5-cre; GOE* mice and stimulated them with (anti-2,4-dinitrophenyl; DNP) IgE and antigen (DNP-coupled human serum albumin) (Starkl et al., 2022). Activated *Gch1*-deficient BMCMCs released significantly decreased serotonin amounts while *Gch1* overexpression resulted in increased serotonin secretion (**Figure 5A**). In line with our *in vivo* observations, altered *Gch1* expression did not influence the amount of secreted histamine nor the pain-inducing cytokine IL-6; moreover, mast cell degranulation responses induced by IgE and antigen or compound 48/80 were unaffected (**Figure S5B**). Of note, the limited maturity of BMCMCs, as compared to *in vivo* connective tissue mast cells, was associated with an incomplete *Mcpt5-cre*-mediated recombination, found only in approximately 35% of cultured BMCMCs (**Figure S5C**) and the extent of BH4 and serotonin reduction in this *in vitro* model may therefore be underestimated. However, these experiments confirm that *Gch1* modulates mast cell BH4 and serotonin synthesis and release while leaving other aspects of mast cell biology and function unaffected.

**Figure 5.**
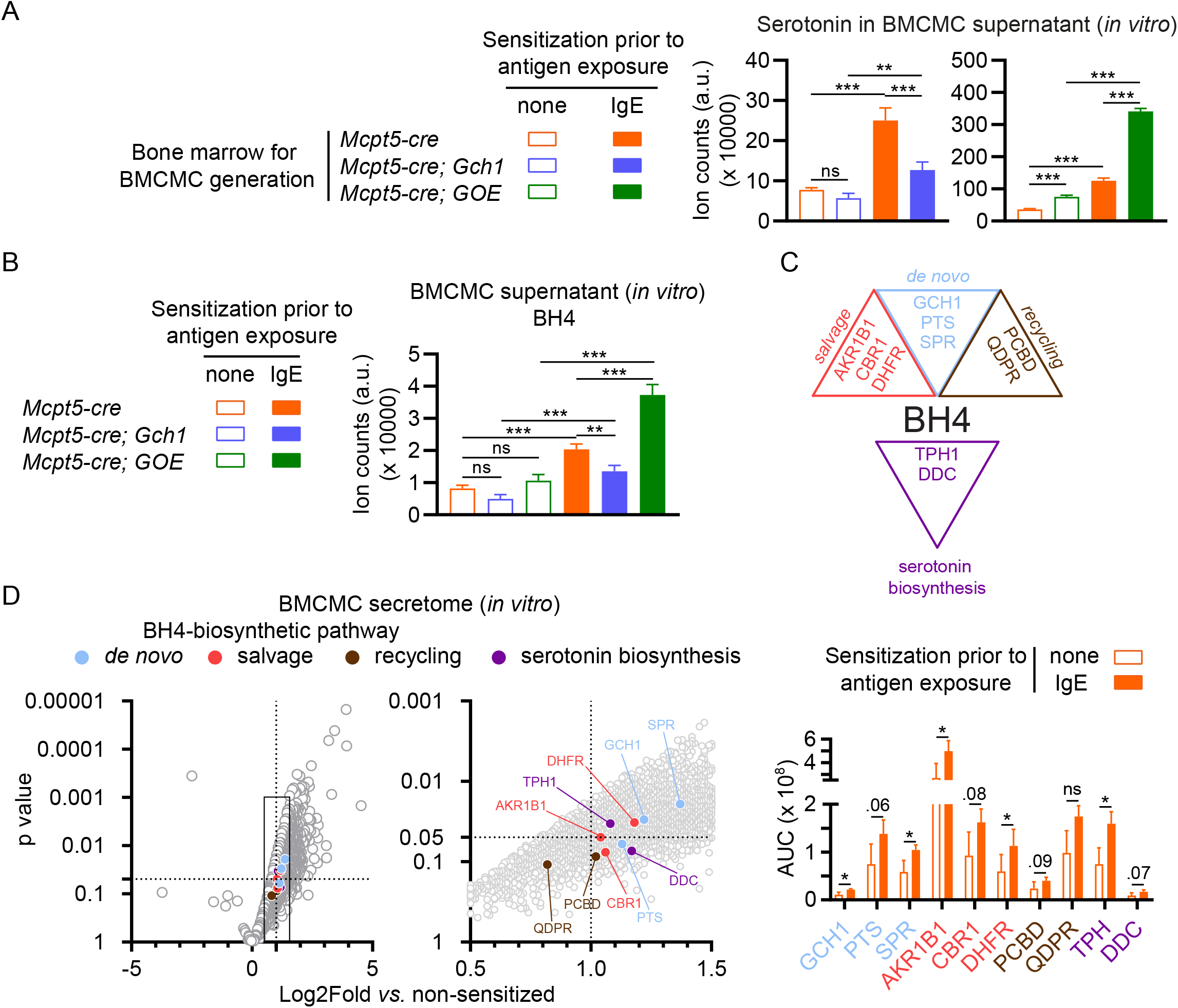
Activated mast cells release BH4, serotonin as well as their biosynthetic enzymes upon degranulation. (A, B) Bone marrow-derived cultured mast cells (BMCMCs) were generated from control (*Mcpt5-cre*) or *Mcpt5-cre; Gch1*, or *Mcpt5-cre; GOE* mice and activated by IgE and antigen for 1 hour, followed by analysis of (A) serotonin or (B) BH4 levels in the supernatant. (C) Schematic depicting the de novo, salvage and recycling arms of the BH4 pathway as well as the serotonin synthesis pathway. GTP, guanosine triphosphate; GCH1, GTP cyclohydrolase I; PTS, 6-Pyruvoyl tetrahydropterin synthase; SPR, sepiapterin reductase; AKR1, aldo-keto reductase family 1; CBR, carbonyl reductase family; DHFR, dihydrofolate reductase; QDPR, quinoid dihydropteridine reductase; PCDB, pterin-4alpha-carbinolamine dehydratase; TPH, tryptophan hydroxylase; DDC, dopa decarboxylase. (D; left and middle panel) Volcano blots of mass spectrometry results representing fold change (x axis) and statistical significance (y axis) of detected proteins in supernatant of IgE-sensitized vs. non-sensitized mast cells after 1 hour antigen exposure (the middle panel is a magnification of the data range indicated by the rectangle in the left panel). Circle colors in the left and middle panels indicate an association with the respective BH4 or serotonin biosynthesis pathways (respective proteins are labelled in the middle panel) as shown in (C) and Figure S5D. The right panel depicts the absolute quantification (area under the curve – AUC) of selected BH4 or serotonin biosynthesis pathways components detected by mass spectrometry. Dotted lines indicated where p=0.05 (horizontal line on the y axis) and a fold change of 2 (vertical line on the x axis). (A, B) One-Way ANOVA with Tukey’s multiple comparisons test; (D, middle and right panels) t-test; * P ≤ 0.05, ** P ≤ 0.01, *** P ≤ 0.001, ns – not significant. Error bars indicate SD. Circular symbols in volcano blots represent detected proteins.

Interestingly, we found that stimulated mast cells not only released serotonin, but also BH4 itself within 1 hour after activation (**Figure 5B**). BH4 can be produced via three (*de novo*, salvage and recycling) major pathways (Werner et al., 2011) (**Figure S5D**). To further investigate the increased release of BH4 and serotonin from degranulated mast cells, we characterized the mast cell secretome by mass spectrometry. Astoundingly, we identified all major enzymes involved in these three BH4 biosynthetic pathways (**Figure 5C and 5D**). Moreover, we identified tryptophan hydroxylase (TPH) and dopa decarboxylase (DDC), the key enzymes necessary for serotonin synthesis from tryptophan (**Figure 5C, 5D and Figure S5D**). Altogether these experiments reveal that mast cells not only release BH4 and serotonin but also the necessary enzymatic machinery required for synthesis of both metabolites rapidly after activation.

### Substance P-mediated hypersensitivity is driven by mast cell BH4

Apart from IgE and antigen-mediated stimulation via the high affinity IgE receptor FcεRI, mast cells respond to a variety of exogenous and endogenous compounds (Redegeld et al., 2018) including the neuropeptide substance P, which potently activates mast cells via a specific receptor, Mas-related GPCR-B2 (Mrgprb2) in mice and MRGPRX2 in humans (Gaudenzio et al., 2016; McNeil et al., 2015). Studies using substance P-deficient mice (Sahbaie et al., 2009) or substance P blocking antibodies (Green et al., 2019) have demonstrated key roles of this neuropeptide in pain hypersensitivity after paw incision. We confirmed that surgical incision leads to a dramatic increase of the substance P precursors α- and β-preprotachykinin in the skin about 8 hours after incision (**Figure 6A**). In contrast to BMCMCs, mouse primary peritoneal mast cells express high levels of Mrgprb2 and respond to substance P (Akula et al., 2020; McNeil et al., 2015). Using enriched peritoneal mast cells (ePMCs) from wild type mice *ex vivo* (**Figure S6A**), we found that substance P induced mast cell degranulation (**Figure S6B**) and the release of serotonin and histamine (**Figure 6B**). Treatment with the Mrgprb2/MRGPRX2 (and neurokinin-1 receptor; NK-1R) tripeptide antagonist QWF (Azimi et al., 2016) significantly blocked the substance P-mediated ePMC serotonin and histamine release (**Figure 6B**).

**Figure 6.**
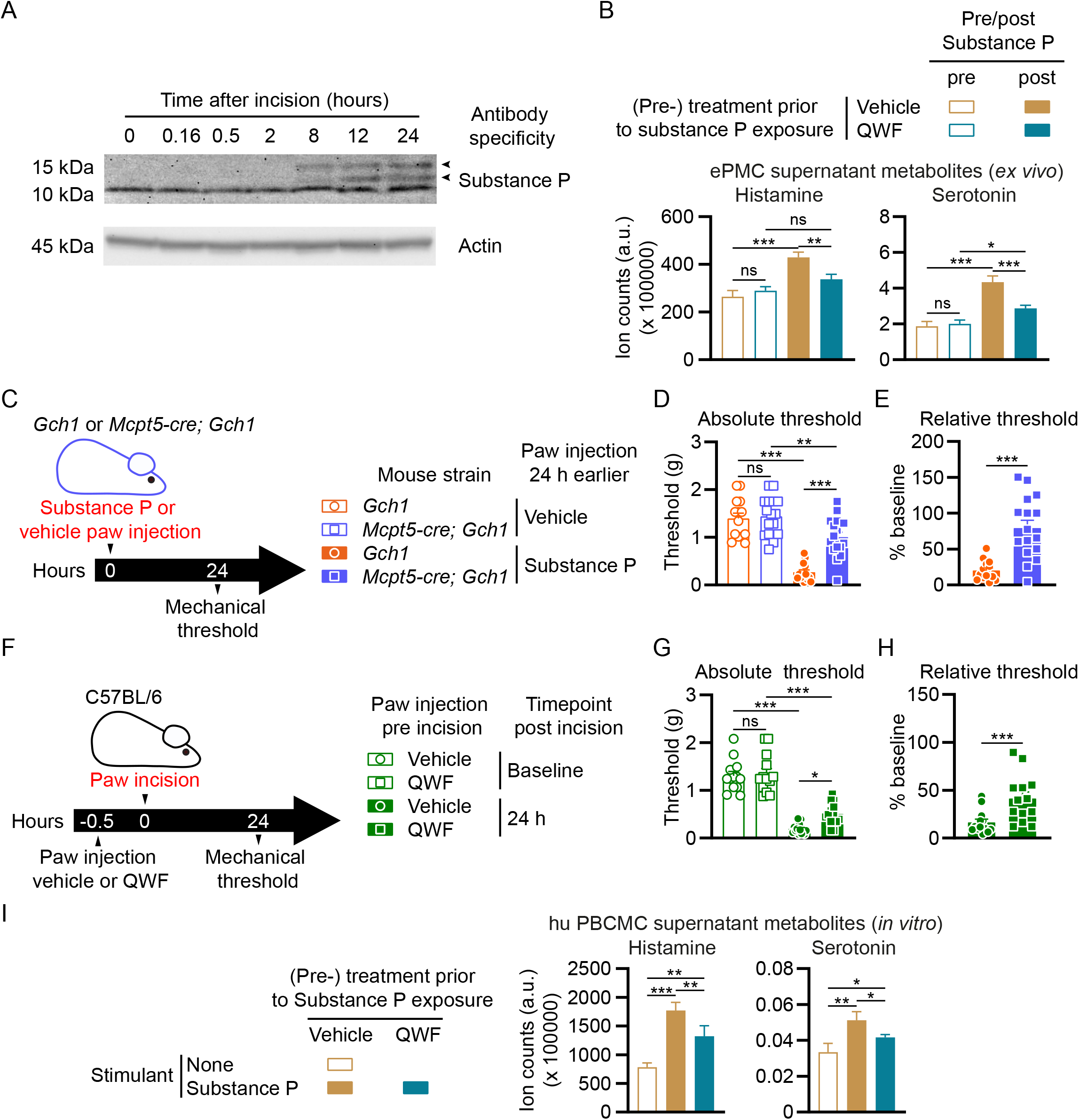
Interference with Substance P-mediated mast cell degranulation alleviates postoperative mechanical hypersensitivity. (A) Western blots of Substance P (upper picture) and actin (lower picture; loading control) in the skin tissue of paws from wild type mice collected before or sampled at the indicated time points after incision. Arrowheads indicate positions of α- and β-preprotachykinin. (B) Enriched peritoneal mast cells (ePMCs) of C57BL/6 wild type mice were pre-treated with vehicle or 100 µM QWF prior to stimulation. Histamine and serotonin in supernatants were analyzed after 1 hour stimulation with 10 µM Substance P. (C) Experimental scheme for (D and E). Control (*Gch1*) and *Mcpt5-cre; Gch1* received injections of either vehicle (saline) or Substance P (50 μg) into the left hind paw. Paw mechanical thresholds were assessed 24 hours after injection. (D) Mechanical thresholds 24 hours post injection (n=14-16). (E) Mechanical thresholds relative to vehicle 24 hours post injection (n=14-16). (F) Experimental scheme for (G and H). C57BL/6 wild type mice were injected with either vehicle or QWF (20 µl of a 0.5mM solution) into the left hind paw 30 minutes before baseline mechanical threshold assessment, followed by paw incision and 24 hours later by mechanical threshold assessment. (G) Absolute mechanical thresholds before and 24 hours post incision (n=13). (H) Mechanical thresholds relative to baseline 24 hours post incision (n=13). (I) Human peripheral blood-derived cultured mast cells (hu PBCMCs) were pre-treated with vehicle or 100 µM QWF prior to simulation (or left unstimulated). Histamine and serotonin in supernatants were analyzed after 1 hour incubation with (or without) 100 µM Substance P. (B, D, G, I) One-Way ANOVA with Tukey’s multiple comparisons test; (E, H) Mann-Whitney test; * P ≤ 0.05, ** P ≤ 0.01, *** P ≤ 0.001, ns – not significant. Error bars indicate (B, I) SD or (D, E, G, H) SEM. Symbols in bar graphs represent individual mice.

Since substance P activates skin mast cells *in vivo* (Gaudenzio et al., 2016) and mast cell-derived BH4 is a critical mediator of postoperative pain (**Figure 3H**), we assessed the possible connection between these observations using *Mcpt5-cre; Gch1* mice (**Figure 6C**). A single intradermal injection of substance P induced pain hypersensitivity after 24 hours in control mice to levels similar to that observed upon paw incision (**Figure 6D and 6E**). Strikingly, substance P-mediated pain hypersensitivity was profoundly blocked in *Mcpt5-cre; Gch1* mice (**Figure 6D and 6E**). To explore the therapeutic application of substance P receptor interference to alleviate postoperative hypersensitivity, we treated wild type mice with QWF prior to surgery (**Figure 6F**). Injecting the compound 30 minutes before paw incision alleviated mechanical allodynia (**Figure 6G and 6H**). QWF treatment of human peripheral blood-derived cultured mast cells (hu PBCMCs; **Figure S6C**) prior to substance P stimulation also significantly reduced their degranulation (**Figure S6D**) as well as histamine and serotonin release (**Figure 6I**). These data highlight the key role of mast cell BH4 in substance P-mediated pain hypersensitivity.

## Discussion

Timely and effective pain control after surgery is crucial as the severity of immediate postoperative pain correlates with the likelihood of transition into chronic pain (Fletcher et al., 2015). Neuroimmune interactions are major orchestrators of immune responses in barrier tissues such as the lung, intestine or skin under physiological and pathophysiological conditions (Schiller et al., 2021). Since inflammatory compounds can directly mediate and shape pain sensitivity (Baral et al., 2019), deciphering the coded signals between nerves and immune cells offers a novel avenue to therapeutic interventions for effective pain management. One metabolite that acts in neurons and immune cells is BH4, primarily generated by the enzyme GCH1. BH4 in sensory neurons has been demonstrated to regulate pain thresholds and has a prominent functional role in immune cells such as macrophages and T cells (Cronin et al., 2018; McNeill et al., 2018; Nagarkoti et al., 2019).

By both mining public gene expression datasets and using *Gch1*-GFP reporter mice, we identified neutrophils, macrophages and mast cells as major *Gch1*-expressing immune cells in injured skin. While neutrophils were the most numerous *Gch1*-expressing immune cells after surgical skin incision, genetic ablation of or *Gch1* deletion specifically in neutrophils neither altered pain hypersensitivity nor BH4 levels in the injured paw. Neutrophils do not, therefore, produce BH4 in injured skin or significantly contribute to the associated pain hypersensitivity. Similarly, macrophage-specific *Gch1* deletion did not affect the increased BH4 or pain sensitivity in the skin after incision. The GCH1/BH4 pathway is linked to peripheral sensory neurons in the context of chronic pain in both animal models and human patients (Belfer et al., 2015; Fujita et al., 2020; Latremoliere et al., 2015; Nasser and Moller, 2014; Tegeder et al., 2006). Whereas chronic nerve injury induces *Gch1* expression in injured sensory DRG neurons, we failed to detect *Gch1* induction or *de novo* BH4 synthesis in sensory neurons upon acute skin incision. Moreover, while surgery-induced injury triggered local sensory neuron release of preformed BH4, our data show that the sensory neuron-derived BH4 does not contribute to the postoperative serotonin production and pain hypersensitivity. The function of the sensory neuron-derived BH4 remains to be investigated but our data highlight that the specific cellular source of BH4 is an important indicator for pain threshold regulation in this model.

A striking new finding is the key importance of mast cell GCH1 and BH4 in the pain response to surgical skin injury. Mast cells are resident in most organs and are especially prominent in barrier tissues (St John et al., 2022). The broad expression of various receptors for exogenous and endogenous molecules combined with their capacity of releasing a plethora of mediators has positioned mast cells as central communication hubs between the nervous, immune, vascular, epithelial, and endocrine systems (Mukai et al., 2018; Redegeld et al., 2018; St John et al., 2022). Mast cell-derived molecules can activate nociceptors and contribute to visceral pain sensitivity (Aguilera-Lizarraga et al., 2021; Barbara et al., 2007), intracranial headaches such as migraines (Levy et al., 2007), or neuropathic pain development after nerve injury (Zuo et al., 2003). While previous reports link mast cell activation and degranulation to postoperative pain, most of these studies involved use of chemicals to manipulate mast cell function such as compound 48/80 and sodium cromoglycate (cromolyn) to induce or prevent mast cell degranulation (Oliveira et al., 2011; Yasuda et al., 2013). However, the conclusions from these studies are limited since compound 48/80 can also directly activate neurons (Schemann et al., 2012) and cromolyn exhibits only minor mast cell inhibitory capacity in mice (Oka et al., 2012) but rather affects other immune cells directly, such as neutrophils and macrophages (Holian et al., 1991; Kay et al., 1987). More recently, a genetic approach using DTR expression driven by the *Mrgprb2* promoter combined with DT application showed the importance of *Mrgprb2*-expressing cells in postoperative pain (Green et al., 2019). Using *Mcpt5-cre; DTA* mice which lack connective tissue mast cells (Dudeck et al., 2011; Galli et al., 2020; St John et al., 2022), we now directly show the critical contribution of mast cells to postoperative pain hypersensitivity. Our results further show that the prominent role of mast cells depends on BH4 production since the phenotype of mast cell-deficient mice was recapitulated in *Mcpt5-cre; Gch1* animals. Moreover, specific *Gch1* overexpression in mast cells increased pain-like behavior, further highlighting the importance of mast cell-derived BH4 in promoting pain hypersensitivity after surgical tissue damage.

Activated mast cells release a variety of mediators with pain modulating roles (Mukai et al., 2018). For instance, IL-6, histamine and serotonin enhance nociception after tissue damage in models of inflammatory and neuropathic pain (Medhurst et al., 2008; Wei et al., 2005; Zeitz et al., 2002; Zuo et al., 2003). However, mast cell-specific BH4 modulation had no apparent effect on paw histamine levels after incision or in an *in vitro* BMCMC degranulation assay. Given that BH4 is required for NO synthesis (as a cofactor for NO synthase) and its association with pain (Aley et al., 1998; Freire et al., 2009), we also investigated whether cultured mast cells release NO upon IgE and antigen-mediated activation but were unable to detect quantifiable NO levels (data not shown). IL-6 secretion was also comparable between control, *Gch1*-deficient as well as *Gch1*-overexpressing mast cells. However, regulation of *Gch1* expression and the resulting alterations in BH4 levels markedly affected total serotonin production and the amount of serotonin released by activated mast cells. BH4 is an essential cofactor for tryptophan hydroxylase, the enzyme required for serotonin biosynthesis (Werner et al., 2011). Moreover, the serotoninergic receptors 5-HT3 and 5-HT2A are expressed on nociceptors (Van Steenwinckel et al., 2009; Zeitz et al., 2002). Since decreased paw BH4 levels and the amelioration of hypersensitivity correlated with reduced serotonin levels, we propose a major role for mast cell-derived serotonin in acute tissue injury pain sensitivity. Intriguingly, we also observed that activated mast cells release BH4 *in vitro*. It is possible that such mast cell-released BH4 itself contributes to pain hypersensitivity by direct action on nociceptors (Nasser et al., 2015). We analyzed supernatant of activated mast cells by mass spectrometry and identified every major enzyme of the three BH4 synthesis pathways in addition to serotonin biosynthetic enzymes among the ∼4000 detected proteins (Werner et al., 2011). This intriguing observation raises the possibility that mast cell granules contain the necessary biosynthetic machinery for extracellular BH4 and serotonin synthesis after degranulation.

Although mast cell BH4 has been shown to regulate histamine release and itch responses in mice (Zschiebsch et al., 2019), we did not observe any effect of BH4 on histamine levels or its release upon degranulation. One reason for this discrepancy could be that the cre line, used in the itch study (LysM-cre) to decipher mast cell-specific contributions, appears to be inactive in mast cells (Abram et al., 2014). Research on pain, including postoperative pain, has historically excluded females despite apparent sex differences related to clinical, experimental and treatment aspects (Fillingim et al., 2009). For instance, peripheral serotonin levels are affected by female hormones and the estrous cycle, and this may contribute to the altered pain sensitivities in females (Kaur et al., 2018). Our study indicates that BH4 in mast cells exert similar effects on postoperative pain in both sexes.

One strategy to therapeutically target mast cells to alleviate pain hypersensitivity would be interfering with the GCH1/BH4 pathway to specifically deplete serotonin in these cells. Systemic pharmacological BH4 blockage ameliorates incision-induced pain in mice (Arai et al., 2020) which could be attributed to multiple target points in the pain circuit. We show here that a single local intradermal paw administration of the sepiapterin reductase blocker QM385 significantly decreased BH4 and serotonin levels in the injured skin area. However, this approach did not significantly reduce mechanical allodynia, probably because the drug was not applied continuously. Continuous topical application could be more effective but requires further research.

Substance P directly acts on various immune cell populations *in vivo*, thus linking sensory neurons to immune responses (Perner et al., 2020; Serhan et al., 2019; Talbot et al., 2015; Talbot et al., 2016). Substance P also enhances pain sensitization after incision injury (Green et al., 2019; Sahbaie et al., 2009). We now show that skin incision leads to rapid generation of the substance P precursors α- and β-preprotachykinin (Nawa et al., 1983) and that blocking substance P’s interaction with its receptor Mrgprb2 using the Mrgprb2/MRBPRX2/NK-1R-antagonist QWF (Azimi et al., 2016), markedly inhibits mast cell degranulation and serotonin release from mouse and human mast cells and significantly reduces mechanical allodynia following surgical injury.

In conclusion, we have identified a crucial role of the mast cell BH4 pathway in acute postoperative pain. Our study highlights the therapeutic potential of interfering with the substance P-mast cell axis and mast cell-specific BH4 production, for postoperative pain management, which specifically targets serotonin-related effects while maintaining other functions of activated mast cells.

## Acknowledgments

Metabolomics was performed by the VBCF Metabolomics Facility which is funded by the City of Vienna through the Vienna Business Agency. We thank Axel Roers (Technical University Dresden) for generously providing *Mcpt5*-cre mice. This work benefitted from data assembled by the *ImmGen* consortium (Heng et al., 2008).

## Author contributions

Shane JF Cronin and Philipp Starkl conceived the study with input from Josef Penninger. Shane JF Cronin and Philipp Starkl performed and designed the experiments with help from coauthors as follows: Gustav Johnsson with flow cytometry; Tyler Artner with gene expression analysis; Bruna Lenfer Turnes, Clifford Woolf and Victoria Klang with *in vivo* QM385 testing; Nadine Serhan, Laura-Marie Gail, Georg Stary and Nicolas Gaudenzio with mast cell *in vitro* experiments and reagents; Keith Channon and Sylvia Knapp with critical reagents; Tiago Oliveira and Karel Stejskal with mass spectrometry; Thomas Köcher with metabolomics; Shane JF Cronin, Philipp Starkl and Josef Penninger wrote the paper with input from all authors.

## Material and Methods

### Ethics

All animal experiments were performed in accordance with institutional policies and federal guidelines. The Austrian Federal Ministry of Education, Science and Research approved the corresponding proposals GZ BMBWF-66.015/0033-V/3b/2019 and their amendments for the experiments performed in this study.

### Mice

If not indicated otherwise, 10-16 weeks old mice were used throughout this study. *Mcpt5-cre* (*Tg(Cma1-cre)ARoer*) mice (Scholten et al., 2008) were generously provided by Axel Roers, Technical University Dresden. *MRP8-cre* (*TG(S100A8-cre,-EGFP)1Ilw*; JAX stock #021614) (Passegue et al., 2004), *LysM-cre* (Imai et al., 2008), iDTR (*Gt(ROSA)26Sortm1(HBEGF)Awai*; JAX stock #007900) (Buch et al., 2005), DTA^fl^ (*Gt(ROSA)26Sortm1(DTA)Lky*; JAX stock #009669) (Voehringer et al., 2008) and lox-stop-lox (lsl)-YFP (*Gt(ROSA)26Sortm1(EYFP)Cos*; JAX stock #006148) (Srinivas et al., 2001) mice were originally obtained from the Jackson Labs. Tamoxifen-inducible sensory neuron-specific *Brn3A-cre*^*ERT*^ mice (Shacham-Silverberg et al., 2018), mice with a Cre-dependent *GCH1*-hemagglutinin overexpression cassette to induce BH4 overproduction (Tg(CAG-GCH1)#Wlf; *Gch1*^*OE*^) (Latremoliere et al., 2015) and *Gch1*^*flox*^ mice (targeting exons 2 and 3 of the GCH1 active site) (Chuaiphichai et al., 2014) have previously been described. Mice expressing eGFP under the *Gch1* promoter (*Tg(Gch1-EGFP)GU68Gsat*) were used to label *Gch1*-expressing cells (Latremoliere et al., 2015). Primary (wild type) cells for *in vitro* experiments were derived from C57BL/6J mice bred and housed at the Core Facility Laboratory Animal Breeding and Husbandry of the Medical University of Vienna. Tamoxifen (Sigma, T5648) was first dissolved in ethanol, then diluted 1:20 in corn oil, heated at 55°C and administered intraperitoneally (i.p.; 2 mg/mouse) daily for five consecutive days to induce deletion (Latremoliere et al., 2015).

### Mouse bone marrow-derived cultured mast cells

Bone marrow-derived cultured mast cells (BMCMCs) were generated by culture of bone marrow in DMEM medium (Sigma), supplemented with 10 % fetal calf serum (FCS; Sigma), 1000 U Penicillin/Streptomycin, 1 mM Sodium Pyruvate (both from Gibco), 10 ng/ml recombinant stem cell factor (SCF) and 10 ng/ml IL-3 (both from Peprotech). Femur and tibia bone marrows from one mouse were initially seeded in 10 ml medium in a 25 cm^2^ tissue culture flask (Corning) and transferred after overnight culture to a 75 cm^2^ flask with 5 ml fresh medium. The cells received 5 ml fresh medium twice a week and were transferred to a new flask once a week. Cells were used after at least 6 weeks of culture and a purity of ≥ 95 %.

### Enrichment of mouse peritoneal mast cells

Peritoneal lavages of female C57BL/6 mice were performed with 5 ml lavage buffer, consisting of filter-sterilized (0.2 µm; StarLab) endotoxin-free phosphate buffered saline (PBS; without Ca^2+^ or Mg^2+^; Sigma) containing 1 % bovine serum albumin (BSA; Sigma) and 1 mM EDTA. Lavages of individual mice were pooled and centrifuged 10 min at 500 g, 4°C. Cells were next resuspended in 50 µl (per mouse) lavage buffer containing 0.5 µg biotinylated anti-mouse FcεRIα (clone MAR-1; Biolegend) and incubated 20 minutes at 4°C. After washing and resuspension in 500 µl lavage buffer, cell suspension was mixed with 15 µl (per mouse) washed magnetic beads (Dynabeads MyONE streptavidin T1; Invitrogen). After 20 minutes incubation on ice (and mixing by pipetting every 5 min), bead-bound cells were enriched using an EasySep Magnet (StemCell Technologies) and washed 3 times with lavage buffer and 1 time with RPMI DTT buffer (see below). Bead-bound cells (enriched peritoneal mast cells; ePMCs) were resuspended in RPMI DTT, counted and aliquoted for downstream assays. For metabolite measurements, ePMCs were aliquoted at 10^5^ cells/well (in 12.5 µl) in a 96 well round bottom plate, followed by addition of 12.5 µl RPMI DTT containing 200 µM QWF (an MRGPRX2 and NK_1_ receptor antagonist) or vehicle and incubation at 37°C for 10 min. After addition of 25 µl of 20 µM Substance P in RPMI DTT, cells were incubated at 37°C for 1 hour. The plate was then put on a 96 Well Magnet Plate (Permagen) and the supernatant was transferred to BioPur 1.5 ml microcentrifuge tubes (Eppendorf), frozen in liquid N_2_ and stored at -80°C until downstream analysis.

### Human mast cell culture

Human peripheral blood-derived cultured mast cells (hu PBCMCs) were generated as previously described (Gaudenzio et al., 2016): In brief, buffy coats of healthy blood donors (from the Etablissement Français du Sang, Toulouse, France) were processed for the isolation of CD34^+^ precursor cells using the EasySep Human CD34 Positive Selection Kit (Stemcell Technologies). Isolated cells were cultured for 1-2 weeks in StemSpan Medium (Stemcell Technologies) supplemented with 10 ng/ml interleukin (IL)-3, 50 ng/ml IL-6 (all cytokines from Preprotech), 1% Penicillin/Streptomycin and 3% supernatant of Chinese hamster ovary transfectants secreting SCF. After two weeks, cells were transferred for approximately 10 weeks to IMDM Glutamax supplemented with 1 mM sodium pyruvate, 50 mM ß-mercaptoethanol, insulin-transferrin selenium (all from Thermo Fisher Scientific), 0.5% bovine serum albumin (Sigma), 10 µg/ml ciprofloxacin (Sigma), 50 ng/ml IL-6 (Preprotech) and 3% supernatant of Chinese hamster ovary transfectants secreting SCF. Cells were then tested functionally for degranulation.

### Postoperative pain model

The postoperative pain model was adapted to that previously described (Cowie and Stucky, 2019), Briefly, mice were anesthetized with a ketasol (10 mg/ml) and xylasol (2 mg/ml) injection administered at 100 ul/10 g body weight intraperitoneally. The left hind paw was exposed and firmly positioned, and a 6 mm longitudinal incision was made through the skin of the plantar foot using a #11 lance blade and scalpel. The incision was started 2 mm from the edge of the heel and extended toward the toes. The skin was opened on either side gently by extending the forceps. The underlying plantar flexor digitorum brevis muscle was gently opened by forcep spreading. The skin was closed with two single sutures of 7-0 nylon, and the wound site was covered with antibiotic ointment. The operated paw was designated the ipsi-lateral (Ipsi) paw while the other paw, the right paw, designated the contra-lateral (Contra) paw. After surgery, mice were allowed to recover on heated pads before being returned to their home cage.

### Spared nerve injury (SNI) model

SNI surgery was performed under 3% induction / 2% maintenance with isoflurane on adult mice (8 to 12 weeks old). The tibial and common peroneal branches of the sciatic nerve were tightly ligated with a 5.0 silk suture and transected distally, while the sural nerve was left intact (Decosterd and Woolf, 2000). After injury, the incision was closed, and mice were allowed to recover on heated pads before being returned to their home cage.

### Behavioral phenotyping

All behavioral experiments were conducted in accordance with the Austrian Animal Care and Use Committee guidelines and in a blinded fashion in a quiet room (temperature 22±1°C) from 9 AM to 6 PM. Both sexes were analyzed in this study. Mice were housed with their littermates (2 to 6 mice per cage based on the litters) with food and water *ad libitum*. All animals were maintained under the same conditions (22±1°C, 50% relative humidity, 12-hour light/dark cycle). For behavioral experiments involving transgenic mice, randomization was achieved through the breeding: at the time of weaning mice were separated based on their sex and placed in their new home cage. Only cages with a mixed representation of transgenic mice and their littermates were used for behavioral experiments. All experiments used at least 2 independent litters and were duplicated.

### Determination of pain responses/thresholds

Mice were transferred to the behavioral phenotyping room at least 1 week prior to experiments. Before each experiment, mice were allowed to habituate to the experimental environment for at least 30 minutes prior to any testing.

Mechanical threshold testing (von Frey filaments). Before testing, each mouse was habituated in a small plastic (7.5 × 7.5 × 15 cm) cage for 1 hour. A logarithmic series of calibrated von Frey monofilaments (Stoelting, Wood Dale, IL, United States), with bending forces that ranged from 0.02 to 1.4 g, were applied with the up–down paradigm (Chaplan et al., 1994), starting with the 0.6 g filament. Filaments were applied twice for 2–3 s, with between-application intervals of at least 30 s to avoid sensitization to the mechanical stimuli. The response to the filament was considered positive if immediate licking or biting, flinching or rapid withdrawal of the stimulated paw was observed. The injured area from the incision-conducted paw as well as its equivalent area on the contralateral paw was used as described previously (Cowie and Stucky, 2019).

To determine thermal sensitivity, contact heat pain (also known as hot plate) testing was conducted: mice were placed on a metallic plate heated to a set temperature (50°C) within an acrylic container (Bioseb, France), and the latency for flinching and/or licking one of the hind paws was recorded. Mice were tested before and 24 hours after incision surgery (performed on both hind paws).

### Histology and imaging

Toluidine blue staining of paraffin skin sections was performed as previously described (Starkl et al., 2016). For immunostaining of DRG tissue and hind paw skin, L3-L4 contralateral or ipsilateral DRG tissue from the paw (either after incision or spared nerve injury) or hind paw skin were extracted, fixed in 4% paraformaldehyde dissolved in PBS, cryoprotected in 30% sucrose, and frozen in OCT (Tissue-Tek). Ten-μm thick cryosections were blocked with 1% bovine serum albumin (Sigma-Aldrich)/ 0.1%Triton X-100 in 0.1 M phosphate buffered saline (PBS) and then incubated with primary antibodies overnight at 4°C. After 3 washes in PBS for 10 minutes each, sections were incubated with secondary antibody for 1 hour at room temperature, washed 3 times in PBS (10 minutes each) and mounted using Dako mounting medium (S3023; Agilent). Primary antibodies and detection reagents used: polyclonal chicken anti-GFP (Aves labs); anti-beta-tubulin III (clone 2G7D4; Abcam); Avidin-FITC (Sigma). Secondary antibodies used: Alexa Fluor 488 anti-rabbit (Jackson Immunoresearch laboratories), 1:500; Alexa Fluor 555 anti-mouse (Jackson Immunoresearch laboratories), 1:500 and Alexa Fluor 594 anti-chicken, (Jackson Immunoresearch laboratories), 1:500. All images were assembled for publication using Fiji. For comparative analysis, fluorescence intensity, exposure time, and other parameters were consistent for all conditions in the same experiment.

### Targeted metabolomics

Metabolites were extracted from frozen tissue or cell pellets using a MeOH:ACN:0.5% dithiothreitol (DTT) in water (2:2:1, v/v) (MeOH= Methanol; ACN=Acetonitrile) ice-cold solvent mixture by adding 300 μL of the solvent to excised paw tissue in an screw-lock 2ml tube homogenized with beads for 30 s, incubated in liquid nitrogen for 1 min, followed by vigorous vortex shaking during 2 min. Cell pellets were disrupted by vigorous vortexing and pipetting up/down. Cycles of liquid nitrogen freezing, and tissue/cell homogenizing were repeated for three times. Samples were then centrifugated at 4,000 × g for 10 min at 4 °C. The supernatant was collected and frozen until LC-MS/MS analysis. Reversed phase liquid chromatography-tandem mass spectrometry (LC-MS/MS) was used for the quantification of BH4, sepiapterin, histamine, neopterin and serotonin. Briefly, 1 µl of the extract was injected on a RSLC ultimate 3000 (Thermo Fisher Scientific) directly coupled to a TSQ Vantage mass spectrometer (Thermo Fisher Scientific) via electrospray ionization. A Kinetex C18 column was used (100 Å, 150 × 2.1 mm) at a flow rate of 80 µl/min. LC-MS/MS was performed by employing the selected reaction monitoring (SRM) mode of the instrument using the transitions (quantifiers) 242.1 *m/z* → 166.1 *m/z* (BH4); 238.1 *m/z* → 192.1 *m/z* (sepiapterin); 112.1 *m/z* → 95.1 *m/z* (histamine); 254.1 *m/z* → 206.1 *m/z* (neopterin) and 177.1 *m/z* → 192.1 *m/z* (serotonin) in the positive ion mode. A 7-minute-long linear gradient was used, staring from 100% A (1 % acetonitrile, 0.1 % formic acid in water) to 80% B (0.1 % formic acid in acetonitrile). Freshly prepared DTT (final concentration at 1 mg/ml) was used for stabilizing BH4. Authentic metabolite standards (Merck) were used for determining the optimal collision energies for LC-MS/MS and for validating experimental retention times. The total intensity of the ion counts of a specific transition is proportional to the amount of that metabolite.

### Mass spectrometry

Cell-free mast cell supernatant was collected into protein LoBind tubes after centrifugation (500 *xg*, 5 min, 4°C), and snap-frozen using liquid nitrogen. Samples were stored at -80°C until Liquid Chromatography coupled to Mass Spectrometry (LC-MS/MS) analysis. Supernatants were individually processed using the iST Sample Preparation Kit (PreOmics) with minor modifications to the provided kit protocol. Briefly, to facilitate lysis, and to reduce and alkylate supernatant proteins, 70 µL of iST 2-fold LYSE buffer were added to 70 µL of each sample and tubes were incubated at 95°C, 1000 rpm for 10 min. After 2 min cooling, samples were sonicated twice for 30 sec using a low intensity setting in a water bath sonicator (30 sec rest on ice). Twenty-five µL of the provided DIGEST solution containing Trypsin and LysC were added to perform the protease digestion overnight on a thermoblock (37°C, 500 rpm). Resulting peptides were cleaned and dried under vacuum at 45°C, resuspended in 0.1% Trifluoroacetic acid and stored at -20°C until further use. LC-MS/MS analyses of the supernatants were performed using a Vanquish Neo UHPLC nano-LC system coupled to an Orbitrap Exploris 480 MS system equipped with NG EasySpray ion source (all from Thermo Fischer Scientific). Proteolytic peptides were loaded onto a PepMap Acclaim C18, 5 mm × 300 μm ID, 5 μm particles, 100 Å pore size trap column (Thermo Fisher Scientific) in combined loading mode, with trap column flush direction backward. After loading, the trap column was switched in-line with a double nanoViper PepMap Acclaim C18, 500 mm × 75 μm ID, 2 μm, 100 Å analytical column (Thermo Fisher Scientific) in a Butterfly heater (PST-BPH-20, Phoenix S&T) and operated at 30°C. The analytical column was directly connected to PepSep sprayer 1 (Bruker) equipped with a 10 μm ID fused silica electrospray emitter with an integrated liquid junction (Bruker). Peptides were eluted using a flow rate of 230 nl/min, starting with the mobile phases 98% A (0.1% v/v formic acid in water) and 2% B (80% acetonitrile, 0.1% v/v formic acid) and linearly increasing to 35% B over the next 180 minutes. The column was then equilibrated in combined control mode with equilibration factor 3. The Orbitrap Exploris 480 was operated in data-dependent mode performing full MS scan (m/z range 350-1200, resolution 60000, normalized AGC target=100%), at 3 compensation voltages (CV -45, -60 and -75), each followed by data-dependent MS/MS scans of the most abundant ions to fill 0.8 sec, 1 sec, and 0.8 sec cycle times, respectively. MS/MS spectra were acquired in the positive mode, using a normalized HCD collision energy of 30, isolation width of 1.0 m/z, resolution of 15000, and normalized Automatic Gain Control (AGC) target was set to 100%. The threshold to select precursor ions for MS/MS was set to 2.5× 10^4^. Precursor ions selected for fragmentation (include charge state 2-6) were excluded for 40 sec. The monoisotopic precursor selection (MIPS) mode was set to Peptide and the include isotopes feature was unselected.

### Mass spectrometry data analysis

For peptide identification, the RAW data files were loaded into Proteome Discoverer (version 2.5.0.400, Thermo Scientific). All MS/MS spectra were searched using MSAmanda v2.0.0.19924 (Dorfer et al., 2014). The peptide and fragment mass tolerance was set to ±10 ppm, the maximal number of missed cleavages was set to 2, using tryptic enzymatic specificity without proline restriction. Peptide and protein identification was performed in two steps: for the initial search, the RAW-files were searched against the Uniprot mouse reference database (21963 sequences; 11728051 residues), supplemented with common contaminants. The result was filtered to 1 % FDR on protein using the Percolator algorithm (Kall et al., 2007) integrated in Proteome Discoverer. A sub-database of proteins identified in this search was generated for further processing. For the second step, the RAW-files were searched against the created sub-database using the same settings as above plus considering additional variable modifications: carbamidomethylation of cysteine was set as a fixed modification; oxidation of methionine, phosphorylation on serine, threonine and tyrosine, deamidation on asparagine and glutamine, pyro-glutamine from glutamine on peptide N-terminal, acetylation on protein N-terminus were set as variable modifications. The localization of the post-translational modification sites within the peptides was performed with the tool ptmRS, based on the tool phosphoRS (Taus et al., 2011). Identified hits were filtered again to 1 % FDR at both protein and PSM level. Additionally, an MSAmanda score cut-off at the PSM level of 150 (at least) was applied. Peptides were subjected to label-free quantification using IMP-apQuant (Doblmann et al., 2019).

Proteins were quantified by summing unique and razor peptides and applying intensity-based absolute quantification (iBAQ, (Schwanhausser et al., 2011)). Proteins were filtered to be identified by a minimum of 2 quantified peptides in at least one of the analysed samples. Protein-abundances-normalization was done based on the MaxLFQ algorithm (Cox et al., 2014).

### Flow cytometry

Ipsilateral and contralateral paw skin was harvested 24 hours after incision of the *Gch1*-GFP reporter mice. After harvest, the paw skin was dissociated using scissors and subsequently incubated with 2 mg/ml collagenase IV (ThermoFisher) and 0.2 mg/ml deoxyribonuclease I (ThermoFisher) in RPMI medium for 45 min at 37°C with shaking. Following digestion, the cells were passed through a 70 μm filter and washed with FACS buffer (PBS, 2% FCS). Cells were stained with the viability dye eFluor780 (eBioscience) in PBS at 4°C for 20 minutes and washed with FACS buffer. The Fc receptors of immune cells were blocked with CD16/CD32 Fc block (BD Bioscience) in FACS buffer at 4°C for 10 minutes, followed by direct addition of flow cytometry antibodies for 20 min at 4°C (see Table 1 for complete antibody list). The samples were washed in FACS buffer and immediately acquired using an LSR Fortessa flow cytometer (BD). The data was analyzed using the FlowJo software v10.7.

**Table 1.**
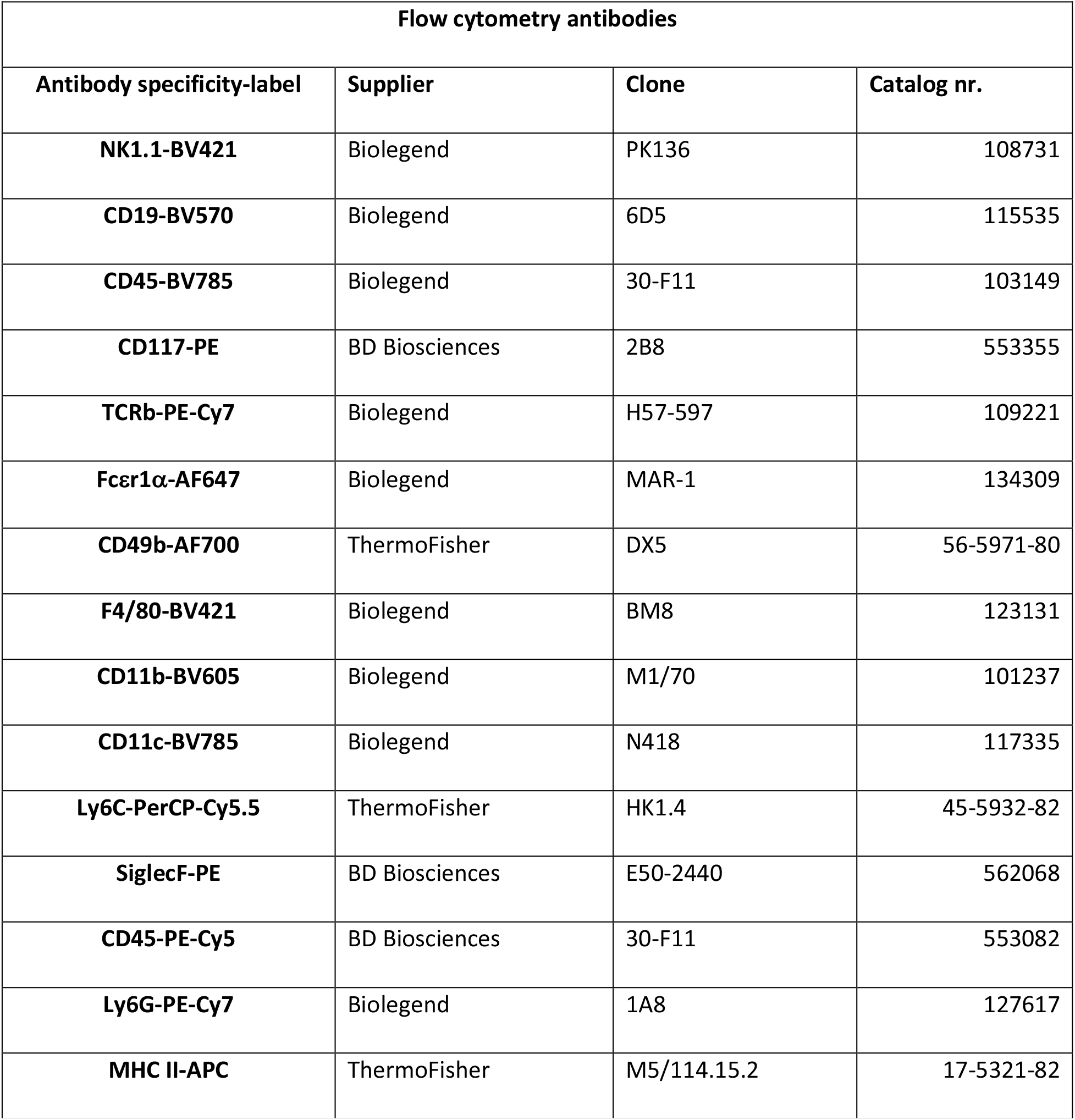
Antibodies used for flow cytometry staining

### *Gch1-GFP* reporter expression analysis

Ipsilateral or contralateral paw skins of C57BL/6 wild type or *Gch1*-GFP, respectively, were collected 24 hours after incision, pooled and processed to generate single cell suspensions and analyzed by flow cytometry as described above. To select potential *Gch1*-expressing candidate immune cell populations in ipsilateral paws of *Gch1*-GFP mice, the GFP mean fluorescence intensities (MFIs) of skin immune cell populations in ipsilateral paws (collected 24 hours after incision injury) of wild type mice (background) was subtracted from the respective MFIs of *Gch1*-GFP animals (see **Figure S1D**, upper panel for the GFP MFI of *Gch1*-GFP immune cell populations). Based on gates excluding background GFP fluorescence signals of respective cell populations analyzed in C57BL/6 wild type mice, cell numbers were determined and populations with high auto fluorescence (i.e. signal equal or lower compared to wild type cells) were excluded from subsequent analysis (see **Figure S1D**, middle panel). Due to high background signal, CD4 T cells, NK cells and monocytes did not pass this first threshold. In a next step, cell populations representing less than 1% of *Gch1*-GFP^+^ leukocytes (dendritic cells, monocytes, eosinophils, B cells, basophils) were excluded (see **Figure S1D**, lower panel).

### Gene expression database/analysis

BioGPS database analysis: *Gch1* (ID 14528) expression in mouse immune cells was analyzed using the BioGPS platform (https://biogps.org) (Wu et al., 2009), Dataset GeneAtlas MOE340 and gcrma, Probesets 1420499_at and 1429692_s_at. Expression values were downloaded and transformed into a heat map using GraphPad Prism v.9.1.

For ImmGen database analysis: For *Gch1* gene expression screening in primary mouse immune cells analyzed by the Immunological Genome Project (ImmGen) (Heng et al., 2008), microarray data were downloaded from the Gene Expression Omnibus (GEO) database (Gene Expression Omnibus ID GSE15907 & GSE37448; see **Table S1** for further details) and post hoc normalized (to all Mouse Gene 1.0 ST Arrays uploaded to the GEO platform) with the platform Gene Expression Commons (GExC) (Seita et al., 2012). Immune cell populations (biological duplicates to quadruplicates) were sorted from male C57BL/6J mice following a standardized procedure (see protocols tab, https://www.immgen.org/) (Dwyer et al., 2016; Ericson et al., 2014; Gautier et al., 2012; Heng et al., 2008; Miller et al., 2012; Robinette et al., 2015). Normalized *Gch1* expression (−100 to 100) profiles were exported from the GExC platform and transformed into a heat map using GraphPad Prism v.9.1.

### Mouse and human mast cell activation and sample collection for metabolomics

For analysis of metabolites in the supernatant of IgE/antigen-activated mouse BMCMCs, cells were sensitized (or not) overnight with 1 µg/ml dinitrophenyl (DNP)-specific IgE (kindly provided by Dr. Fu-Tong Liu (University of California-Davis) (Liu et al., 1980). Next day, cells were collected and washed twice in RPMI 1640 without phenol red (Gibco) containing 1 mg/ml DTT (Roche; RPMI DTT). Cells were resuspended at 2× 10^6^ cells/ml in RPMI DTT and seeded at 50 µl aliquots (quadruplicates) in a 96-well round bottom tissue culture plate (Corning). After addition of 50 µl of 20 ng dinitrophenyl_30-40_-conjugated human serum albumin (DNP-HAS; Sigma)/ml in RPMI/DTT, cells were incubated for 1 hour at 37°C. Cells were then transferred to Bio-pur 1.5 ml microcentrifuge tubes (Eppendorf) and centrifuged for 5 min at 500 x g, 4°C. After transfer of supernatant to fresh tubes, supernatant and pellets were frozen in liquid N_2_ and stored at -80°C. IL-6 was measured using the ELISA MAX Standard Set Mouse IL-6 (Biolegend).

For assessment of Substance P-mediated mast cell activation and mediator release, ePMCs or hu PBCMCs (2.5× 10^5^ cells in 50 µl RPMI DTT) were either pre-treated with 100 µM QWF or DMSO for 10 minutes, followed by stimulation with 10 µM or 100 µM Substance P, respectively. After 1 hour incubation, cells were processed as described above for BMCMCs.

### Drugs/reagents/treatments

QM385 has been previously developed by our group (Cronin et al., 2018). For *in vitro* use, both sepiapterin and QM385 were dissolved in DMSO to a stock concentration of 10 mM. Working concentrations are indicated in the figures. For *in vivo* preparation of QM385, the compound was dissolved in 1% Tween 80 with 5% carboxy methylcellulose, medium viscosity (Sigma). The preparation was sonicated for 5 minutes and applied (20 μl of 10 mM solution) intradermally 2 hours before incision surgery. Tamoxifen (Sigma; dissolved in ethanol and corn oil; see above) was administered i.p. to the *Brn3A-Ert-Cre* expressing mice for four consecutive days four weeks before the experiment. Diphtheria toxin (Sigma) was dissolved in water and administered i.p. (25 ng/g body weight) for two days to deplete neutrophils in *MRP8-cre; DTR* mice. Substance P (#1156; dissolved in saline) and Boc-GlnD-Trp(Formyl)-Phe-OBzl (QWF; #6642; dissolved in DMSO) were from Tocris.

### Statistical analysis

Technical replicates were performed in cell-based *in vitro* degranulation and activation experiments and were measured in quadruplicates. Statistical analysis was performed using GraphPad Prism 9.3.1 (GraphPad Software) except for mass spectrometry analysis in which LIMMA was used. All necessary details are stated in the respective figure legends.

## References

Abram, C.L., G.L. Roberge, Y. Hu, and C.A. Lowell. 2014. Comparative analysis of the efficiency and specificity of myeloid-Cre deleting strains using ROSA-EYFP reporter mice. J Immunol Methods 408:89–100.

Aguilera-Lizarraga, J., M.V. Florens, M.F. Viola, P. Jain, L. Decraecker, I. Appeltans, M. Cuende-Estevez, N. Fabre, K. Van Beek, E. Perna, D. Balemans, N. Stakenborg, S. Theofanous, G. Bosmans, S.U. Mondelaers, G. Matteoli, S. Ibiza Martinez, C. Lopez-Lopez, J. Jaramillo-Polanco, K. Talavera, Y.A. Alpizar, T.B. Feyerabend, H.R. Rodewald, R. Farre, F.A. Redegeld, J. Si, J. Raes, C. Breynaert, R. Schrijvers, C. Bosteels, B.N. Lambrecht, S.D. Boyd, R.A. Hoh, D. Cabooter, M. Nelis, P. Augustijns, S. Hendrix, J. Strid, R. Bisschops, D.E. Reed, S.J. Vanner, A. Denadai-Souza, M.M. Wouters, and G.E. Boeckxstaens. 2021. Local immune response to food antigens drives meal-induced abdominal pain. Nature 590:151–156.

Akula, S., A. Paivandy, Z. Fu, M. Thorpe, G. Pejler, and L. Hellman. 2020. How Relevant Are Bone Marrow-Derived Mast Cells (BMMCs) as Models for Tissue Mast Cells? A Comparative Transcriptome Analysis of BMMCs and Peritoneal Mast Cells. Cells 9:

Aley, K.O., G. McCarter, and J.D. Levine. 1998. Nitric oxide signaling in pain and nociceptor sensitization in the rat. J Neurosci 18:7008–7014.

Arai, H., R. Takahashi, Y. Sakamoto, T. Kitano, O. Mashita, S. Hara, S. Yoshikawa, K. Kawasaki, and H. Ichinose. 2020. Peripheral tetrahydrobiopterin is involved in the pathogenesis of mechanical hypersensitivity in a rodent postsurgical pain model. Pain 161:2520–2531.

Arias, J.I., M.A. Aller, and J. Arias. 2009. Surgical inflammation: a pathophysiological rainbow. J Transl Med 7:19.

Azimi, E., V.B. Reddy, K.C. Shade, R.M. Anthony, S. Talbot, P.J.S. Pereira, and E.A. Lerner. 2016. Dual action of neurokinin-1 antagonists on Mas-related GPCRs. JCI Insight 1:e89362.

Baral, P., S. Udit, and I.M. Chiu. 2019. Pain and immunity: implications for host defence. Nat Rev Immunol 19:433–447.

Barbara, G., B. Wang, V. Stanghellini, R. de Giorgio, C. Cremon, G. Di Nardo, M. Trevisani, B. Campi, P. Geppetti, M. Tonini, N.W. Bunnett, D. Grundy, and R. Corinaldesi. 2007. Mast cell-dependent excitation of visceral-nociceptive sensory neurons in irritable bowel syndrome. Gastroenterology 132:26–37.

Belfer, I., F. Dai, H. Kehlet, P. Finelli, L. Qin, R. Bittner, and E.K. Aasvang. 2015. Association of functional variations in COMT and GCH1 genes with postherniotomy pain and related impairment. Pain 156:273–279.

Brennan, T.J., E.P. Vandermeulen, and G.F. Gebhart. 1996. Characterization of a rat model of incisional pain. Pain 64:493–502.

Buch, T., F.L. Heppner, C. Tertilt, T.J. Heinen, M. Kremer, F.T. Wunderlich, S. Jung, and A. Waisman. 2005. A Cre-inducible diphtheria toxin receptor mediates cell lineage ablation after toxin administration. Nat Methods 2:419–426.

Chaplan, S.R., F.W. Bach, J.W. Pogrel, J.M. Chung, and T.L. Yaksh. 1994. Quantitative assessment of tactile allodynia in the rat paw. J Neurosci Methods 53:55–63.

Chuaiphichai, S., E. McNeill, G. Douglas, M.J. Crabtree, J.K. Bendall, A.B. Hale, N.J. Alp, and K.M. Channon. 2014. Cell-autonomous role of endothelial GTP cyclohydrolase 1 and tetrahydrobiopterin in blood pressure regulation. Hypertension 64:530–540.

Cowie, A.M., and C.L. Stucky. 2019. A Mouse Model of Postoperative Pain. Bio Protoc 9:

Cox, J., M.Y. Hein, C.A. Luber, I. Paron, N. Nagaraj, and M. Mann. 2014. Accurate proteome-wide label-free quantification by delayed normalization and maximal peptide ratio extraction, termed MaxLFQ. Mol Cell Proteomics 13:2513–2526.

Cronin, S.J.F., S. Rao, M.A. Tejada, B.L. Turnes, S. Licht-Mayer, T. Omura, C. Brenneis, E. Jacobs, L. Barrett, A. Latremoliere, N. Andrews, K.M. Channon, A. Latini, A.C. Arvanites, L.S. Davidow, M. Costigan, L.L. Rubin, J.M. Penninger, and C.J. Woolf. 2022. Phenotypic drug screen uncovers the metabolic GCH1/BH4 pathway as key regulator of EGFR/KRAS-mediated neuropathic pain and lung cancer. Sci Transl Med 14:eabj1531.

Cronin, S.J.F., C. Seehus, A. Weidinger, S. Talbot, S. Reissig, M. Seifert, Y. Pierson, E. McNeill, M.S. Longhi, B.L. Turnes, T. Kreslavsky, M. Kogler, D. Hoffmann, M. Ticevic, D. da Luz Scheffer, L. Tortola, D. Cikes, A. Jais, M. Rangachari, S. Rao, M. Paolino, M. Novatchkova, M. Aichinger, L. Barrett, A. Latremoliere, G. Wirnsberger, G. Lametschwandtner, M. Busslinger, S. Zicha, A. Latini, S.C. Robson, A. Waisman, N. Andrews, M. Costigan, K.M. Channon, G. Weiss, A.V. Kozlov, M. Tebbe, K. Johnsson, C.J. Woolf, and J.M. Penninger. 2018. The metabolite BH4 controls T cell proliferation in autoimmunity and cancer. Nature 563:564–568.

Decosterd, I., and C.J. Woolf. 2000. Spared nerve injury: an animal model of persistent peripheral neuropathic pain. Pain 87:149–158.

Doblmann, J., F. Dusberger, R. Imre, O. Hudecz, F. Stanek, K. Mechtler, and G. Durnberger. 2019. apQuant: Accurate Label-Free Quantification by Quality Filtering. J Proteome Res 18:535–541.

Dolin, S.J., J.N. Cashman, and J.M. Bland. 2002. Effectiveness of acute postoperative pain management: I. Evidence from published data. Br J Anaesth 89:409–423.

Dorfer, V., P. Pichler, T. Stranzl, J. Stadlmann, T. Taus, S. Winkler, and K. Mechtler. 2014. MS Amanda, a universal identification algorithm optimized for high accuracy tandem mass spectra. J Proteome Res 13:3679–3684.

Dudeck, A., J. Dudeck, J. Scholten, A. Petzold, S. Surianarayanan, A. Kohler, K. Peschke, D. Vohringer, C. Waskow, T. Krieg, W. Muller, A. Waisman, K. Hartmann, M. Gunzer, and A. Roers. 2011. Mast cells are key promoters of contact allergy that mediate the adjuvant effects of haptens. Immunity 34:973–984.

Dwyer, D.F., N.A. Barrett, K.F. Austen, and C. Immunological Genome Project. 2016. Expression profiling of constitutive mast cells reveals a unique identity within the immune system. Nat Immunol 17:878–887.

Ericson, J.A., P. Duffau, K. Yasuda, A. Ortiz-Lopez, K. Rothamel, I.R. Rifkin, P.A. Monach, and C. ImmGen. 2014. Gene expression during the generation and activation of mouse neutrophils: implication of novel functional and regulatory pathways. PLoS One 9:e108553.

Fillingim, R.B., C.D. King, M.C. Ribeiro-Dasilva, B. Rahim-Williams, and J.L. Riley, 3rd. 2009. Sex, gender, and pain: a review of recent clinical and experimental findings. J Pain 10:447–485.

Fletcher, D., U.M. Stamer, E. Pogatzki-Zahn, R. Zaslansky, N.V. Tanase, C. Perruchoud, P. Kranke, M. Komann, T. Lehman, W. Meissner, and C.g.f.t.C.T.N.g.o.t.E.S.o.A. eu. 2015. Chronic postsurgical pain in Europe: An observational study. Eur J Anaesthesiol 32:725–734.

Foley, B.S. 2006. Wall and Melzack’s Textbook of Pain, 5th Edition. American Journal of Physical Medicine & Rehabilitation 85:581.

Freire, M.A., J.S. Guimaraes, W.G. Leal, and A. Pereira. 2009. Pain modulation by nitric oxide in the spinal cord. Front Neurosci 3:175–181.

Fujita, M., D.D.L. Scheffer, B.L. Turnes, S.J.F. Cronin, A. Latremoliere, M. Costigan, C.J. Woolf, A. Latini, and N.A. Andrews. 2020. Sepiapterin Reductase Inhibition Leading to Selective Reduction of Inflammatory Joint Pain in Mice and Increased Urinary Sepiapterin Levels in Humans and Mice. Arthritis Rheumatol 72:57–66.

Galli, S.J., N. Gaudenzio, and M. Tsai. 2020. Mast Cells in Inflammation and Disease: Recent Progress and Ongoing Concerns. Annu Rev Immunol 38:49–77.

Gan, T.J. 2017. Poorly controlled postoperative pain: prevalence, consequences, and prevention. J Pain Res 10:2287–2298.

Gaudenzio, N., R. Sibilano, T. Marichal, P. Starkl, L.L. Reber, N. Cenac, B.D. McNeil, X. Dong, J.D. Hernandez, R. Sagi-Eisenberg, I. Hammel, A. Roers, S. Valitutti, M. Tsai, E. Espinosa, and S.J. Galli. 2016. Different activation signals induce distinct mast cell degranulation strategies. J Clin Invest 126:3981–3998.

Gautier, E.L., T. Shay, J. Miller, M. Greter, C. Jakubzick, S. Ivanov, J. Helft, A. Chow, K.G. Elpek, S. Gordonov, A.R. Mazloom, A. Ma’ayan, W.J. Chua, T.H. Hansen, S.J. Turley, M. Merad, G.J. Randolph, and C. Immunological Genome. 2012. Gene-expression profiles and transcriptional regulatory pathways that underlie the identity and diversity of mouse tissue macrophages. Nat Immunol 13:1118–1128.

Ghasemlou, N., I.M. Chiu, J.P. Julien, and C.J. Woolf. 2015. CD11b+Ly6G-myeloid cells mediate mechanical inflammatory pain hypersensitivity. Proc Natl Acad Sci U S A 112:E6808–6817.

Green, D.P., N. Limjunyawong, N. Gour, P. Pundir, and X. Dong. 2019. A Mast-Cell-Specific Receptor Mediates Neurogenic Inflammation and Pain. Neuron 101:412–420 e413.

Heng, T.S., M.W. Painter, and C. Immunological Genome Project. 2008. The Immunological Genome Project: networks of gene expression in immune cells. Nat Immunol 9:1091–1094.

Hill, C.E., B.J. Harrison, K.K. Rau, M.T. Hougland, M.B. Bunge, L.M. Mendell, and J.C. Petruska. 2010. Skin incision induces expression of axonal regeneration-related genes in adult rat spinal sensory neurons. J Pain 11:1066–1073.

Holian, A., R. Hamilton, and R.K. Scheule. 1991. Mechanistic aspects of cromolyn sodium action on the alveolar macrophage: inhibition of stimulation by soluble agonists. Agents Actions 33:318–325.

Imai, Y., K. Kuba, G.G. Neely, R. Yaghubian-Malhami, T. Perkmann, G. van Loo, M. Ermolaeva, R. Veldhuizen, Y.H. Leung, H. Wang, H. Liu, Y. Sun, M. Pasparakis, M. Kopf, C. Mech, S. Bavari, J.S. Peiris, A.S. Slutsky, S. Akira, M. Hultqvist, R. Holmdahl, J. Nicholls, C. Jiang, C.J. Binder, and J.M. Penninger. 2008. Identification of oxidative stress and Toll-like receptor 4 signaling as a key pathway of acute lung injury. Cell 133:235–249.

Kall, L., J.D. Canterbury, J. Weston, W.S. Noble, and M.J. MacCoss. 2007. Semi-supervised learning for peptide identification from shotgun proteomics datasets. Nat Methods 4:923–925.

Kaur, S., W.L. Benton, S.A. Tongkhuya, C.M.C. Lopez, L. Uphouse, and D.L. Averitt. 2018. Sex Differences and Estrous Cycle Effects of Peripheral Serotonin-Evoked Rodent Pain Behaviors. Neuroscience 384:87–100.

Kay, A.B., G.M. Walsh, R. Moqbel, A.J. MacDonald, T. Nagakura, M.P. Carroll, and H.B. Richerson. 1987. Disodium cromoglycate inhibits activation of human inflammatory cells in vitro. J Allergy Clin Immunol 80:1–8.

Kehlet, H., T.S. Jensen, and C.J. Woolf. 2006. Persistent postsurgical pain: risk factors and prevention. Lancet 367:1618–1625.

Latremoliere, A., A. Latini, N. Andrews, S.J. Cronin, M. Fujita, K. Gorska, R. Hovius, C. Romero, S. Chuaiphichai, M. Painter, G. Miracca, O. Babaniyi, A.P. Remor, K. Duong, P. Riva, L.B. Barrett, N. Ferreiros, A. Naylor, J.M. Penninger, I. Tegeder, J. Zhong, J. Blagg, K.M. Channon, K. Johnsson, M. Costigan, and C.J. Woolf. 2015. Reduction of Neuropathic and Inflammatory Pain through Inhibition of the Tetrahydrobiopterin Pathway. Neuron 86:1393–1406.

Levy, D., R. Burstein, V. Kainz, M. Jakubowski, and A.M. Strassman. 2007. Mast cell degranulation activates a pain pathway underlying migraine headache. Pain 130:166–176.

Liu, F.T., J.W. Bohn, E.L. Ferry, H. Yamamoto, C.A. Molinaro, L.A. Sherman, N.R. Klinman, and D.H. Katz. 1980. Monoclonal dinitrophenyl-specific murine IgE antibody: preparation, isolation, and characterization. J Immunol 124:2728–2737.

Liu, Q.Q., X.X. Yao, S.H. Gao, R. Li, B.J. Li, W. Yang, and R.J. Cui. 2020. Role of 5-HT receptors in neuropathic pain: potential therapeutic implications. Pharmacol Res 159:104949.

McNeil, B.D., P. Pundir, S. Meeker, L. Han, B.J. Undem, M. Kulka, and X. Dong. 2015. Identification of a mast-cell-specific receptor crucial for pseudo-allergic drug reactions. Nature 519:237–241.

McNeill, E., E. Stylianou, M.J. Crabtree, R. Harrington-Kandt, A.L. Kolb, M. Diotallevi, A.B. Hale, P. Bettencourt, R. Tanner, M.K. O’Shea, M. Matsumiya, H. Lockstone, J. Muller, H.A. Fletcher, D.R. Greaves, H. McShane, and K.M. Channon. 2018. Regulation of mycobacterial infection by macrophage Gch1 and tetrahydrobiopterin. Nat Commun 9:5409.

Medhurst, S.J., S.D. Collins, A. Billinton, S. Bingham, R.G. Dalziel, A. Brass, J.C. Roberts, A.D. Medhurst, and I.P. Chessell. 2008. Novel histamine H3 receptor antagonists GSK189254 and GSK334429 are efficacious in surgically-induced and virally-induced rat models of neuropathic pain. Pain 138:61–69.

Miller, J.C., B.D. Brown, T. Shay, E.L. Gautier, V. Jojic, A. Cohain, G. Pandey, M. Leboeuf, K.G. Elpek, J. Helft, D. Hashimoto, A. Chow, J. Price, M. Greter, M. Bogunovic, A. Bellemare-Pelletier, P.S. Frenette, G.J. Randolph, S.J. Turley, M. Merad, and C. Immunological Genome. 2012. Deciphering the transcriptional network of the dendritic cell lineage. Nat Immunol 13:888–899.

Mukai, K., M. Tsai, H. Saito, and S.J. Galli. 2018. Mast cells as sources of cytokines, chemokines, and growth factors. Immunol Rev 282:121–150.

Nagarkoti, S., S. Sadaf, D. Awasthi, T. Chandra, K. Jagavelu, S. Kumar, and M. Dikshit. 2019. L-Arginine and tetrahydrobiopterin supported nitric oxide production is crucial for the microbicidal activity of neutrophils. Free Radic Res 53:281–292.

Nasser, A., S. Ali, S. Wilsbech, O.J. Bjerrum, and L.B. Moller. 2015. Intraplantar injection of tetrahydrobiopterin induces nociception in mice. Neurosci Lett 584:247–252.

Nasser, A., and L.B. Moller. 2014. GCH1 variants, tetrahydrobiopterin and their effects on pain sensitivity. Scand J Pain 5:121–128.

Nawa, H., T. Hirose, H. Takashima, S. Inayama, and S. Nakanishi. 1983. Nucleotide sequences of cloned cDNAs for two types of bovine brain substance P precursor. Nature 306:32–36.

Oka, T., J. Kalesnikoff, P. Starkl, M. Tsai, and S.J. Galli. 2012. Evidence questioning cromolyn’s effectiveness and selectivity as a ‘mast cell stabilizer’ in mice. Lab Invest 92:1472–1482.

Oliveira, S.M., C.C. Drewes, C.R. Silva, G. Trevisan, S.L. Boschen, C.G. Moreira, D. de Almeida Cabrini, C. Da Cunha, and J. Ferreira. 2011. Involvement of mast cells in a mouse model of postoperative pain. Eur J Pharmacol 672:88–95.

Passegue, E., E.F. Wagner, and I.L. Weissman. 2004. JunB deficiency leads to a myeloproliferative disorder arising from hematopoietic stem cells. Cell 119:431–443.

Perner, C., C.H. Flayer, X. Zhu, P.A. Aderhold, Z.N.A. Dewan, T. Voisin, R.B. Camire, O.A. Chow, I.M. Chiu, and C.L. Sokol. 2020. Substance P Release by Sensory Neurons Triggers Dendritic Cell Migration and Initiates the Type-2 Immune Response to Allergens. Immunity 53:1063–1077 e1067.

Pogatzki, E.M., and S.N. Raja. 2003. A mouse model of incisional pain. Anesthesiology 99:1023–1027.

Reber, L.L., C.M. Gillis, P. Starkl, F. Jonsson, R. Sibilano, T. Marichal, N. Gaudenzio, M. Berard, S. Rogalla, C.H. Contag, P. Bruhns, and S.J. Galli. 2017. Neutrophil myeloperoxidase diminishes the toxic effects and mortality induced by lipopolysaccharide. J Exp Med 214:1249–1258.

Redegeld, F.A., Y. Yu, S. Kumari, N. Charles, and U. Blank. 2018. Non-IgE mediated mast cell activation. Immunol Rev 282:87–113.

Robinette, M.L., A. Fuchs, V.S. Cortez, J.S. Lee, Y. Wang, S.K. Durum, S. Gilfillan, M. Colonna, and C. Immunological Genome. 2015. Transcriptional programs define molecular characteristics of innate lymphoid cell classes and subsets. Nat Immunol 16:306–317.

Rosa, A.C., and R. Fantozzi. 2013. The role of histamine in neurogenic inflammation. Br J Pharmacol 170:38–45.

Sahbaie, P., X. Shi, T.Z. Guo, Y. Qiao, D.C. Yeomans, W.S. Kingery, and D.J. Clark. 2009. Role of substance P signaling in enhanced nociceptive sensitization and local cytokine production after incision. Pain 145:341–349.

Schemann, M., E.M. Kugler, S. Buhner, C. Eastwood, J. Donovan, W. Jiang, and D. Grundy. 2012. The mast cell degranulator compound 48/80 directly activates neurons. PLoS One 7:e52104.

Schiller, M., T.L. Ben-Shaanan, and A. Rolls. 2021. Neuronal regulation of immunity: why, how and where? Nat Rev Immunol 21:20–36.

Scholten, J., K. Hartmann, A. Gerbaulet, T. Krieg, W. Muller, G. Testa, and A. Roers. 2008. Mast cell-specific Cre/loxP-mediated recombination in vivo. Transgenic Res 17:307–315.

Schwanhausser, B., D. Busse, N. Li, G. Dittmar, J. Schuchhardt, J. Wolf, W. Chen, and M. Selbach. 2011. Global quantification of mammalian gene expression control. Nature 473:337–342.

Segelcke, D., B. Pradier, S. Reichl, L.C. Schafer, and E.M. Pogatzki-Zahn. 2021. Investigating the Role of Ly6G(+) Neutrophils in Incisional and Inflammatory Pain by Multidimensional Pain-Related Behavioral Assessments: Bridging the Translational Gap. Front Pain Res (Lausanne) 2:735838.

Seita, J., D. Sahoo, D.J. Rossi, D. Bhattacharya, T. Serwold, M.A. Inlay, L.I. Ehrlich, J.W. Fathman, D.L. Dill, and I.L. Weissman. 2012. Gene Expression Commons: an open platform for absolute gene expression profiling. PLoS One 7:e40321.

Serhan, N., L. Basso, R. Sibilano, C. Petitfils, J. Meixiong, C. Bonnart, L.L. Reber, T. Marichal, P. Starkl, N. Cenac, X. Dong, M. Tsai, S.J. Galli, and N. Gaudenzio. 2019. House dust mites activate nociceptor-mast cell clusters to drive type 2 skin inflammation. Nat Immunol 20:1435–1443.

Shacham-Silverberg, V., H. Sar Shalom, R. Goldner, Y. Golan-Vaishenker, N. Gurwicz, I. Gokhman, and A. Yaron. 2018. Phosphatidylserine is a marker for axonal debris engulfment but its exposure can be decoupled from degeneration. Cell Death Dis 9:1116.

Shipton, E.A., E.E. Shipton, and A.J. Shipton. 2018. A Review of the Opioid Epidemic: What Do We Do About It? Pain Ther 7:23–36.

Sommer, C. 2004. Serotonin in pain and analgesia: actions in the periphery. Mol Neurobiol 30:117–125.

Srinivas, S., T. Watanabe, C.S. Lin, C.M. William, Y. Tanabe, T.M. Jessell, and F. Costantini. 2001. Cre reporter strains produced by targeted insertion of EYFP and ECFP into the ROSA26 locus. BMC Dev Biol 1:4.

St John, A.L., A.P.S. Rathore, and F. Ginhoux. 2022. New perspectives on the origins and heterogeneity of mast cells. Nat Rev Immunol

Starkl, P., N. Gaudenzio, T. Marichal, L.L. Reber, R. Sibilano, M.L. Watzenboeck, F. Fontaine, A.C. Mueller, M. Tsai, S. Knapp, and S.J. Galli. 2022. IgE antibodies increase honeybee venom responsiveness and detoxification efficiency of mast cells. Allergy 77:499–512.

Starkl, P., T. Marichal, N. Gaudenzio, L.L. Reber, R. Sibilano, M. Tsai, and S.J. Galli. 2016. IgE antibodies, FcepsilonRIalpha, and IgE-mediated local anaphylaxis can limit snake venom toxicity. J Allergy Clin Immunol 137:246–257 e211.

Talbot, S., R.E. Abdulnour, P.R. Burkett, S. Lee, S.J. Cronin, M.A. Pascal, C. Laedermann, S.L. Foster, J.V. Tran, N. Lai, I.M. Chiu, N. Ghasemlou, M. DiBiase, D. Roberson, C. Von Hehn, B. Agac, O. Haworth, H. Seki, J.M. Penninger, V.K. Kuchroo, B.P. Bean, B.D. Levy, and C.J. Woolf. 2015. Silencing Nociceptor Neurons Reduces Allergic Airway Inflammation. Neuron 87:341–354.

Talbot, S., S.L. Foster, and C.J. Woolf. 2016. Neuroimmunity: Physiology and Pathology. Annu Rev Immunol 34:421–447.

Tauber, M., F. Wang, B. Kim, and N. Gaudenzio. 2021. Bidirectional sensory neuron-immune interactions: a new vision in the understanding of allergic inflammation. Curr Opin Immunol 72:79–86.

Taus, T., T. Kocher, P. Pichler, C. Paschke, A. Schmidt, C. Henrich, and K. Mechtler. 2011. Universal and confident phosphorylation site localization using phosphoRS. J Proteome Res 10:5354–5362.

Tegeder, I., M. Costigan, R.S. Griffin, A. Abele, I. Belfer, H. Schmidt, C. Ehnert, J. Nejim, C. Marian, J. Scholz, T. Wu, A. Allchorne, L. Diatchenko, A.M. Binshtok, D. Goldman, J. Adolph, S. Sama, S.J. Atlas, W.A. Carlezon, A. Parsegian, J. Lotsch, R.B. Fillingim, W. Maixner, G. Geisslinger, M.B. Max, and C.J. Woolf. 2006. GTP cyclohydrolase and tetrahydrobiopterin regulate pain sensitivity and persistence. Nat Med 12:1269–1277.

Van Steenwinckel, J., A. Noghero, K. Thibault, M.J. Brisorgueil, J. Fischer, and M. Conrath. 2009. The 5-HT2A receptor is mainly expressed in nociceptive sensory neurons in rat lumbar dorsal root ganglia. Neuroscience 161:838–846.

Voehringer, D., H.E. Liang, and R.M. Locksley. 2008. Homeostasis and effector function of lymphopenia-induced “memory-like” T cells in constitutively T cell-depleted mice. J Immunol 180:4742–4753.

Wei, H., Y. Chen, and Y. Hong. 2005. The contribution of peripheral 5-hydroxytryptamine2A receptor to carrageenan-evoked hyperalgesia, inflammation and spinal Fos protein expression in the rat. Neuroscience 132:1073–1082.

Weiser, T.G., S.E. Regenbogen, K.D. Thompson, A.B. Haynes, S.R. Lipsitz, W.R. Berry, and A.A. Gawande. 2008. An estimation of the global volume of surgery: a modelling strategy based on available data. Lancet 372:139–144.

Werner, E.R., N. Blau, and B. Thony. 2011. Tetrahydrobiopterin: biochemistry and pathophysiology. Biochem J 438:397–414.

Wu, C., C. Orozco, J. Boyer, M. Leglise, J. Goodale, S. Batalov, C.L. Hodge, J. Haase, J. Janes, J.W. Huss, 3rd, and A.I. Su. 2009. BioGPS: an extensible and customizable portal for querying and organizing gene annotation resources. Genome Biol 10:R130.

Yasuda, M., K. Kido, N. Ohtani, and E. Masaki. 2013. Mast cell stabilization promotes antinociceptive effects in a mouse model of postoperative pain. J Pain Res 6:161–166.

Zeitz, K.P., N. Guy, A.B. Malmberg, S. Dirajlal, W.J. Martin, L. Sun, D.W. Bonhaus, C.L. Stucky, D. Julius, and A.I. Basbaum. 2002. The 5-HT3 subtype of serotonin receptor contributes to nociceptive processing via a novel subset of myelinated and unmyelinated nociceptors. J Neurosci 22:1010–1019.

Zschiebsch, K., C. Fischer, A. Wilken-Schmitz, G. Geisslinger, K. Channon, K. Watschinger, and I. Tegeder. 2019. Mast cell tetrahydrobiopterin contributes to itch in mice. J Cell Mol Med 23:985–1000.

Zuo, Y., N.M. Perkins, D.J. Tracey, and C.L. Geczy. 2003. Inflammation and hyperalgesia induced by nerve injury in the rat: a key role of mast cells. Pain 105:467–479.

