## Supplemental figures, table and legends for "Mast cell-derived BH4 is a critical mediator of postoperative pain"

[1] Research Division of Infection Biology, Department of Medicine I, Medical University of Vienna, Vienna, Austria, [2] Institute of Molecular Biotechnology of the Austrian Academy of Sciences, Vienna, Austria, [3] Vienna BioCenter PhD Program, Doctoral School of the University of Vienna and Medical University of Vienna, Vienna, Austria, [4] Department of Internal Medicine II, Medical University of Vienna, Vienna, Austria, [5] Department of Neurobiology, Harvard Medical School, Boston, United States, [6] F.M. Kirby Neurobiology Research Center, Boston Children's Hospital, Boston, United States, Department of Pathology, Medical University of Vienna, Vienna, Austria, [7] Toulouse Institute for Infectious and Inflammatory Diseases (Infinity), Inserm UMR1291 CNRS UMR5051, University of Toulouse III, Toulouse, France, [8] Department of Dermatology, Medical University of Vienna, Vienna, Austria, [9] LBI-RUD – Ludwig-Boltzmann Institute for Rare and Undiagnosed Diseases, Vienna, Austria, [10] CeMM, Research Center for Molecular Medicine of the Austrian Academy of Sciences, Vienna, Austria, [11] Radcliffe Department of, British Heart Foundation Centre of Research Excellence, John Radcliffe Hospital, University of Oxford, Oxford, UK, [12] Vienna BioCenter Core Facilities (VBCF), 1030 Vienna, Austria, [13] Department of Pharmaceutical Sciences, University of Vienna, Vienna, Austria, [14] Genoskin SAS, Toulouse, France, [15] Department of Medical Genetics, Life Sciences Institute, University of British Columbia, Vancouver, Canada

### Author list footnotes

*corresponding author

### Corresponding authors email addresses

Figure S1

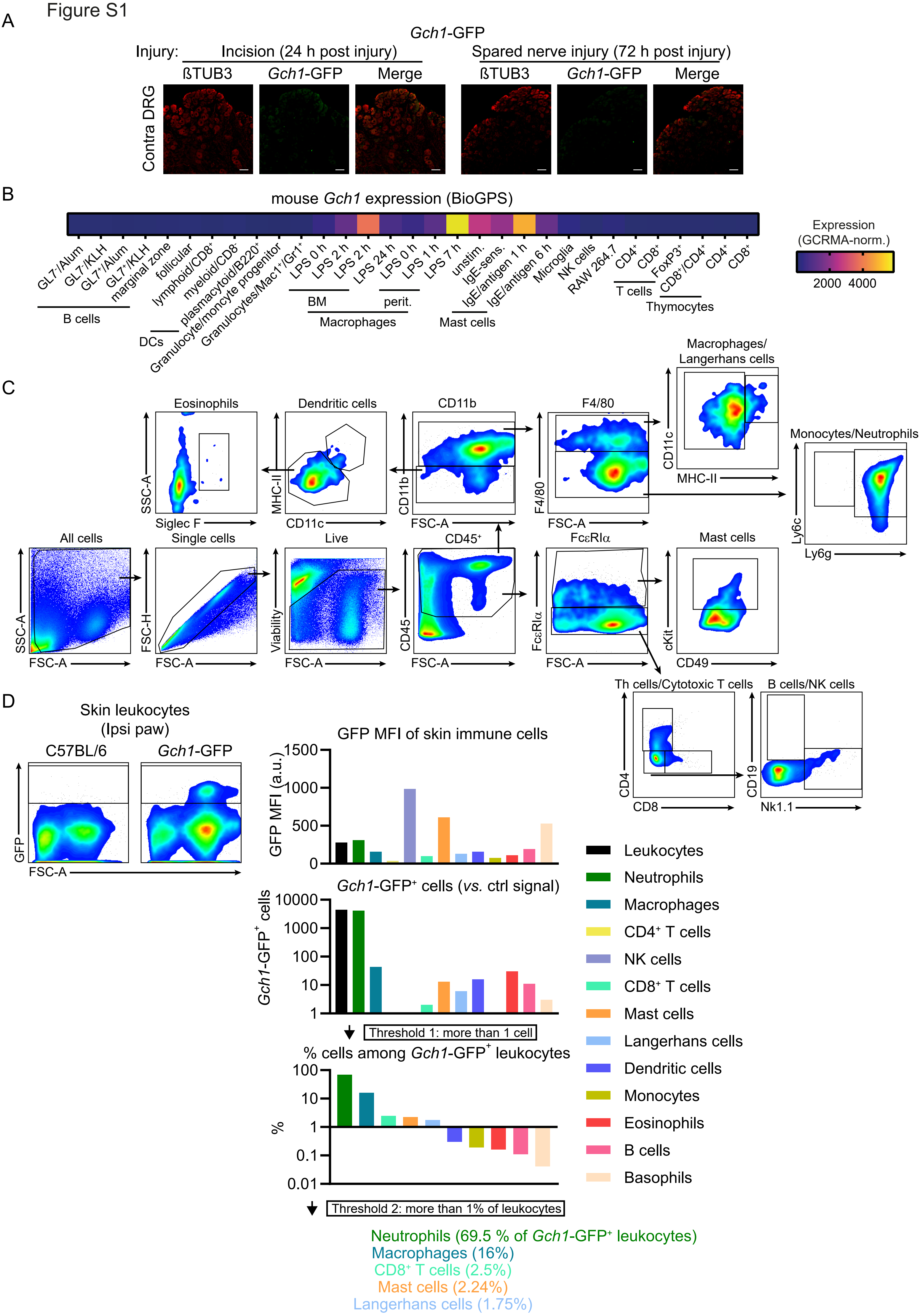
A
Gch1-GFP
Injury:
Incision (24 h post injury)
Spared nerve injury (72 h post injury)
ßTUB3
Gch1-GFP
Merge
Contra DRG
B
mouse Gch1 expression (BioGPS)
GL7-/Alum
GL7-/KLH
GL7+/Alum
GL7+/KLH
marginal zone
follicular
B cells
lymphoid/CD8+
myeloid/CD8-
plasmacytoid/B220+
DCs
Granulocyte/moncyte progenitor
Granulocytes/Mac1+/Gr1+
LPS 0 h
LPS 2 h
LPS 2 h
LPS 24 h
LPS 0 h
LPS 1 h
LPS 7 h
BM
perit.
Macrophages
unstim.
IgE-sens.
IgE/antigen 1 h
IgE/antigen 6 h
Mast cells
Microglia
NK cells
RAW 264.7
CD4+
CD8+
FoxP3+
T cells
CD8+/CD4+
CD4+
CD8+
Thymocytes
Expression
(GCRMA-norm.)
2000
4000
C
Eosinophils
Dendritic cells
CD11b
F4/80
Macrophages/
Langerhans cells
Monocytes/Neutrophils
SSC-A
Siglec F
MHC-II
CD11c
CD11b
FSC-A
F4/80
FSC-A
CD11c
MHC-II
Ly6c
Ly6g
All cells
Single cells
Live
CD45+
FcεRIα
Mast cells
SSC-A
FSC-A
FSC-H
FSC-A
Viability
FSC-A
CD45
FSC-A
FcεRIα
FSC-A
cKit
CD49
Th cells/Cytotoxic T cells
B cells/NK cells
CD4
CD8
CD19
Nk1.1
D
Skin leukocytes
(Ipsi paw)
C57BL/6
Gch1-GFP
GFP
FSC-A
GFP MFI of skin immune cells
GFP MFI (a.u.)
1500
1000
500
0
Gch1-GFP+ cells (vs. ctrl signal)
Gch1-GFP+ cells
10000
1000
100
10
1
Threshold 1: more than 1 cell
% cells among Gch1-GFP+ leukocytes
%
100
10
1
0.1
0.01
Threshold 2: more than 1% of leukocytes
Leukocytes
Neutrophils
Macrophages
CD4+ T cells
NK cells
CD8+ T cells
Mast cells
Langerhans cells
Dendritic cells
Monocytes
Eosinophils
B cells
Basophils
Neutrophils (69.5 % of Gch1-GFP+ leukocytes)
Macrophages (16%)
CD8+ T cells (2.5%)
Mast cells (2.24%)
Langerhans cells (1.75%)

### Figure S1. BioGPS *Gch1* gene expression, flow cytometry gating strategy and selection strategy of *Gch1*-expressing cell populations

(A) Representative immunofluorescence images of anti-beta-TUBULIN-III (ßTUB3) and anti-GFP (*Gch1*-GFP) staining in the contralateral (Contra) L4-L5 dorsal root ganglion tissues of incision injury- (representative of n=3) and spared nerve injury (SNI)-treated (representative of n=3) *Gch1*-GFP reporter mice. Scale bars represent 50 µm. (B) Mean gene expression levels of *Gch1* (n=4) in immune cell populations derived from the *BioGPS* database, Dataset GeneAtlas MOE340, gcrma, Probesets 1420499_at and 1429692_s_at. (C) Gating strategy for identification of skin immune cell populations. Data shown are from skin samples of ipsilateral paws collected from mice 24 hours after incision injury. All leukocyte populations are subgates of live/CD45$^+$ cells; Dendritic cells: CD11b$^-$/CD11c$^+$/MHCII$^+$; Mast cells: Fc$\varepsilon$RI$\alpha^+$/cKit$^+$; Eosinophils: CD11b$^-$/MHCII$^-$/Siglec F$^+$/SSC-A$^{high}$; Macrophages: CD11b$^+$/F4/80$^+$/MHCII$^-$; Langerhans cells: CD11b$^+$/F4/80$^+$/CD11c$^+$/MHCII$^+$; Monocytes: CD11b$^+$/F4/80$^-$/Ly6c$^+$/Ly6g$^-$; Neutrophils: CD11b$^+$/F4/80$^-$/Ly6c$^+$/Ly6g$^+$; Th cells: Fc$\varepsilon$RI$\alpha^-$/CD4$^+$/CD8$^-$/; Cytotoxic T cells: Fc$\varepsilon$RI$\alpha^-$/CD8$^+$/CD4$^-$/; NK cells: Fc$\varepsilon$RI$\alpha^-$/CD19$^-$/CD8$^-$/CD4$^-$/NK1.1$^+$; B cells: Fc$\varepsilon$RI$\alpha^-$/CD8$^-$/CD4$^-$/NK1.1$^-$/CD19$^+$; (D) Selection strategy of *Gch1*-expressing candidate immune cell populations in ipsilateral paws of *Gch1*-GFP mice 24 hours after incision injury (shown in Figure 2C). The upper panel shows GFP mean fluorescence intensities (MFIs) of indicated skin immune cell populations in ipsilateral paws (collected 24 hours after incision injury) of *Gch1*-GFP mice after subtraction of respective GFP MFIs of C57BL/6 wild type mice. The middle panel shows the number of *Gch1*-GFP$^+$ cells among the respective immune cell populations defined by gates excluding background GFP fluorescence signals based respective cell populations analyzed in C57BL/6 wild type mice. Of note, no CD4 T cells, NK cells and monocytes with signal above background were detected and therefore were excluded from the subsequent analysis step. The lower panel shows the proportions of the indicated cell populations (excluding CD4 T cells, NK cells and monocytes) among *Gch1*-GFP$^+$ leukocytes (smoothed dot plots show GFP signals of skin CD45$^+$ leukocytes of C57BL/6 *vs*. *Gch1*-GFP ipsilateral paws). Cell populations representing less than 1% of *Gch1*-GFP$^+$ leukocytes (dendritic cells, monocytes, eosinophils, B cells, basophils) were neglected as potential candidates. Those cell types meeting set arbitrary thresholds (more than 1 cell with GFP signal above background; cell population represents more than 1 % of *Gch1*-GFP$^+$ leukocytes), namely neutrophils, macrophages, CD8$^+$ T cells, mast cells and Langerhans cells are represented in Figure 2C.

Figure S2

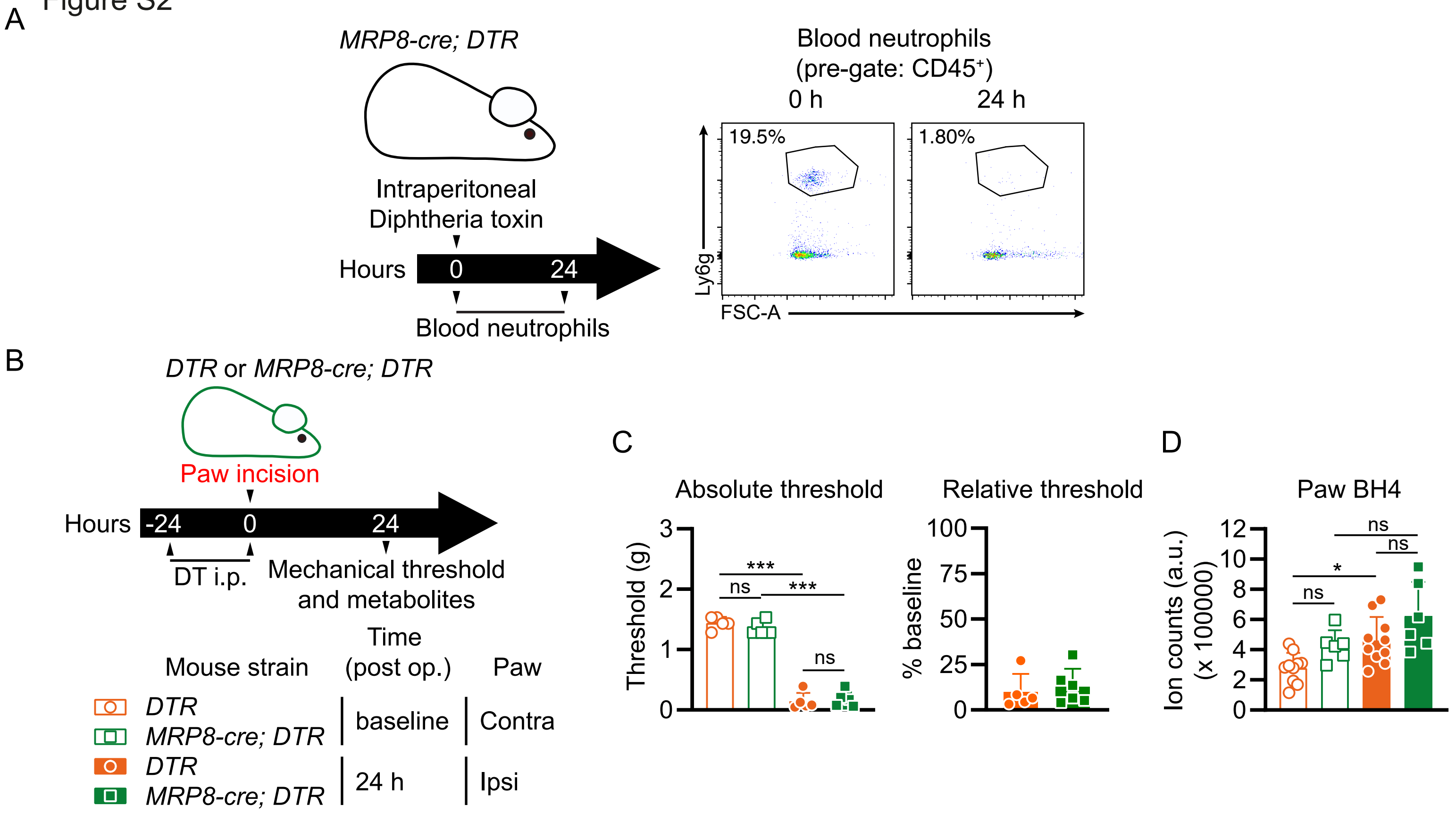
A
MRP8-cre; DTR
Intraperitoneal
Diphtheria toxin
Hours
0
24
Blood neutrophils
Blood neutrophils
(pre-gate: CD45+)
0 h
24 h
19.5%
1.80%
Ly6g
FSC-A
B
DTR or MRP8-cre; DTR
Paw incision
Hours
-24
0
24
DT i.p.
Mechanical threshold
and metabolites
Mouse strain
Time
(post op.)
Paw
DTR
MRP8-cre; DTR
baseline
Contra
DTR
MRP8-cre; DTR
24 h
Ipsi
C
Absolute threshold
Threshold (g)
***
ns
***
ns
Relative threshold
% baseline
D
Paw BH4
Ion counts (a.u.)
(x 100000)
ns
ns
*
ns

### Figure S2. Depletion efficiency, mechanical pain threshold and paw BH4 levels of neutrophil-deficient mice

(A) After blood collection for assessment of baseline neutrophil levels, *MRP8-cre; DTR* mice (which express diphtheria toxin receptor specifically on neutrophils) were treated by intraperitoneal injection of diphtheria toxin (DT) and blood neutrophil numbers analyzed by flow cytometry 24 hours later. Dot plots show samples collected from the same mouse. Percentages refer to the proporation of (gated) Ly6g+ cells selected as indicated by the gates. (B) Experimental scheme for C and D. Control (*DTR*) and *MRP8-cre; DTR* mice were injected intraperitoneally (i.p.) with diphtheria toxin (DT) 24 hours before and at the time of incision. Contralateral (Contra) and ipsilateral (Ipsi) paw mechanical pain sensitivity and BH4 levels were assessed 24 hours after incision. (C) Absolute mechanical (left panel) threshold at baseline and 24 hours post incision and (right panel) thresholds relative to baseline (n=5-6). (D) Paw BH4 levels (n=6-10).

(C, D) One-Way ANOVA with Tukey's multiple comparisons test; * $P \le 0.05$, *** $P \le 0.001$, ns – not significant. (C, D) Error bars indicate SD. Symbols in bar graphs represent individual mice.

Figure S3

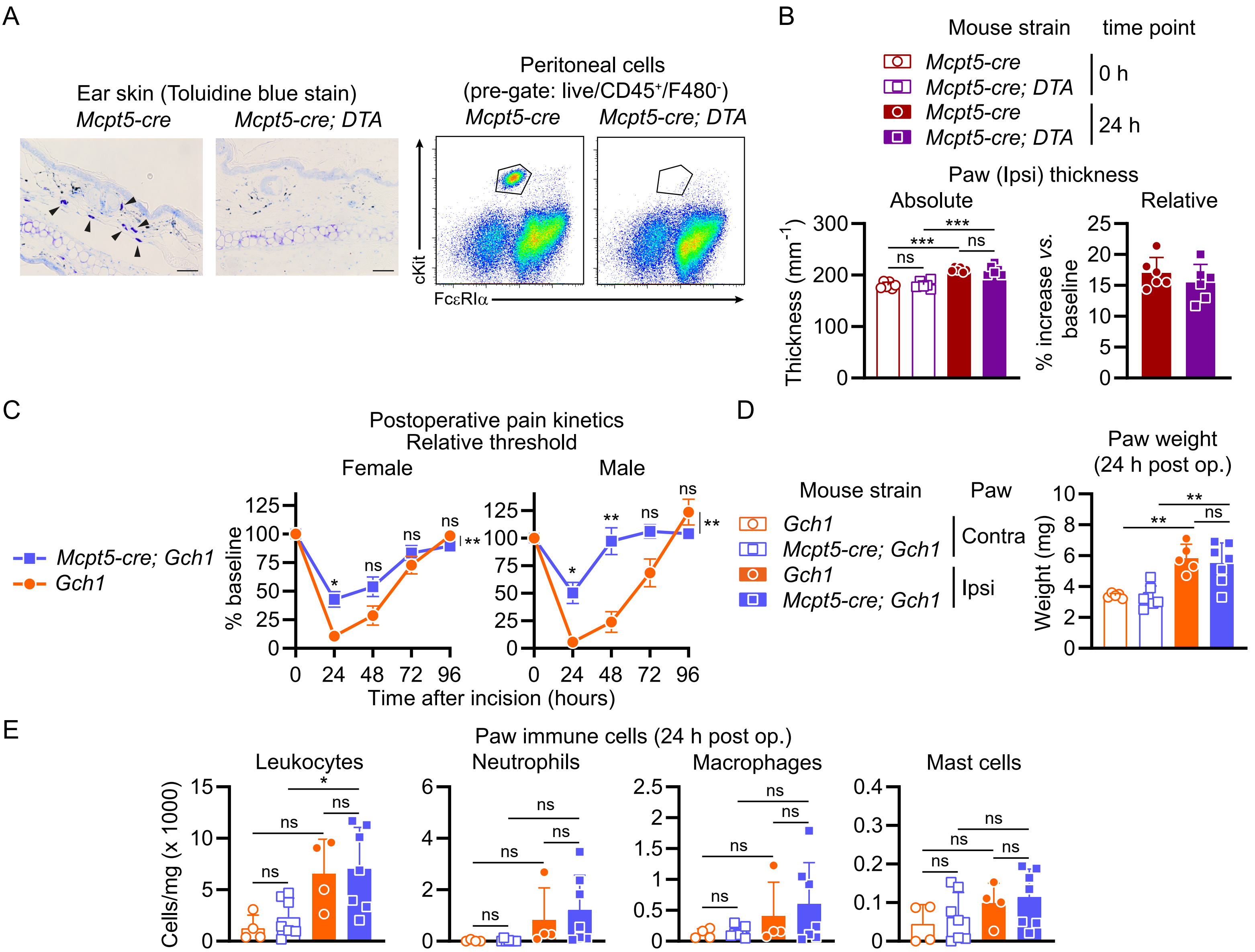

A
Ear skin (Toluidine blue stain)
Mcpt5-cre
Mcpt5-cre; DTA
Peritoneal cells
(pre-gate: live/CD45+/F480-)
Mcpt5-cre
Mcpt5-cre; DTA
cKit
FcεRIα
B
Mouse strain
time point
Mcpt5-cre
Mcpt5-cre; DTA
0 h
Mcpt5-cre
Mcpt5-cre; DTA
24 h
Paw (Ipsi) thickness
Absolute
Relative
Thickness (mm-1)
% increase vs. baseline
C
Postoperative pain kinetics
Relative threshold
Female
Male
Mcpt5-cre; Gch1
Gch1
% baseline
Time after incision (hours)
D
Mouse strain
Paw
Gch1
Mcpt5-cre; Gch1
Contra
Gch1
Mcpt5-cre; Gch1
Ipsi
Paw weight
(24 h post op.)
Weight (mg)
E
Paw immune cells (24 h post op.)
Leukocytes
Neutrophils
Macrophages
Mast cells
Cells/mg (x 1000)

### Figure S3. Mast cell deficiency and paw thickness of *Mcpt5-cre; Gch1* mice and additional physiological and immunological parameters of *Mcpt5-cre; Gch1* mice after incision injury

(A) Left panel: Toluidine-blue stained ear skin tissue sections of *Mcpt5-cre* (control) and *Mcpt5-cre; DTA* (mast cell-deficient) mice. Arrowheads indicate tissue mast cells in control mice. Scale bars indicate 50 µm. Right panel: flow cytometry dot plots showing mast cell gates (cKit$^+$/Fc$\varepsilon$RI$\alpha^+$) among peritoneal lavage cells derived from control and mast cell-deficient mice. (B) Absolute and relative (to baseline) thickness of Ipsi paws of *Mcpt5-cre* and *Mcpt5-cre; DTA* mice 24 hours after incision injury (n=6). (C to E) Postoperative pain kinetics, paw weight and paw skin immune cells were assessed in male and female control (*Gch1*) and *Mcpt5-cre; Gch1* mice. (C) Time course of relative (to baseline) mechanical thresholds of female and male mice at indicated timepoints after incision (n=5-8). (D) Weight of Contra and Ipsi paws 24 hours after incision (n=5-7). (E) Skin immune cell populations in Contra and Ipsi paws 24 hours after incision (n=4-7). (B, D, E) One-Way ANOVA with Tukey's multiple comparisons test; (C) Two-Way ANOVA with Sidak's multiple comparisons test (comparing individual time points) and Two-Way ANOVA with repeated measures with Geisser-Greenhouse correction (overall comparison); * P ≤ 0.05, ** P ≤ 0.01, *** P ≤ 0.001, ns – not significant. Error bars indicate (B, D, E) SD or (C) SEM. Symbols in bar graphs represent individual mice.

Figure S4

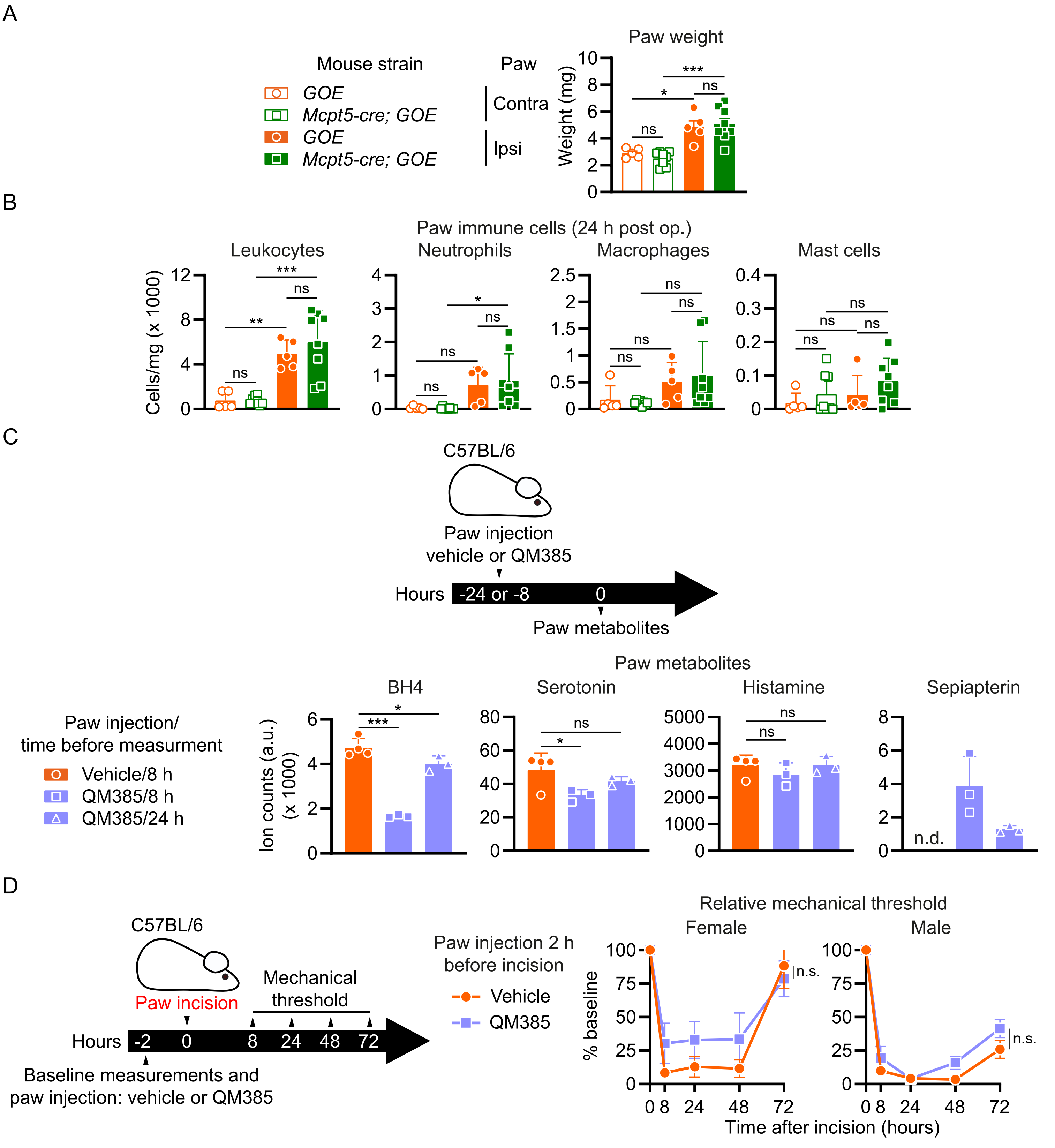
A
Mouse strain
GOE
Mcpt5-cre; GOE
GOE
Mcpt5-cre; GOE
Paw
Contra
Ipsi
Paw weight
Weight (mg)
ns
*
***
ns
B
Paw immune cells (24 h post op.)
Leukocytes
Neutrophils
Macrophages
Mast cells
Cells/mg (x 1000)
ns
**
***
*
C
C57BL/6
Paw injection
vehicle or QM385
Hours
-24 or -8
0
Paw metabolites
Paw metabolites
BH4
Serotonin
Histamine
Sepiapterin
Paw injection/
time before measurment
Vehicle/8 h
QM385/8 h
QM385/24 h
Ion counts (a.u.)
(x 1000)
n.d.
D
C57BL/6
Paw incision
Mechanical
threshold
Hours
-2
0
8
24
48
72
Baseline measurements and
paw injection: vehicle or QM385
Paw injection 2 h
before incision
Vehicle
QM385
Relative mechanical threshold
Female
Male
% baseline
n.s.
Time after incision (hours)

### Figure S4. Postoperative paw weight and skin immune cells of mice with mast cell-specific *Gch1* overexpression and effects of sepipaterin reductase inhibitor (QM385) treatment on paw metabolites and incision-induced postoperative pain hypersensitivity

(A) Paw weight and (B) skin immune cells of contralateral (Contra) and ipsilateral (Ipsi) paws in control (*GOE*) mice and animals with mast cell-specific *Gch1*-overexpression (*Mcpt5-cre; GOE*) 24 hours after incision injury. (C) Upper panel: experimental outline; Left hindpaws of C57BL/6 wild type mice were injected intradermally with either vehicle or QM385, a sepiapterin reductase inhibitor, at 24 or 8 hours before measurement of metabolites in the skin. Lower panels: Paw metabolite levels 8 (vehicle and QM385) or 24 hours (QM385) after compound injection (n=3-4). (D) Left panel: experimental outline; Left hindpaws of C57BL/6 wild type mice were injected with either vehicle or QM385 at 2 hours before incision injury, followed by measurement of relative (to baseline) mechanical threshold kinetics. Right panels: Pain thresholds kinetics of female (n=10) and male (n=8-13) mice (which had received vehicle or QM385 injections in the paws 2 hours before incision) at indicated time points after incision.

(A, B and C) One-Way ANOVA with (A and B) Tukey's or (C) Dunnett's multiple comparisons test; * $P \leq 0.05$, ** $P \leq 0.01$, *** $P \leq 0.001$, ns – not significant. (D) Two-Way ANOVA with Sidak's multiple comparisons test (comparing individual time points) and Two-Way ANOVA with repeated measures with Geisser-Greenhouse correction (overall comparison); Error bars indicate ( and C) SD or (A and D) SEM. Symbols in bar graphs represent individual mice.

Figure S5

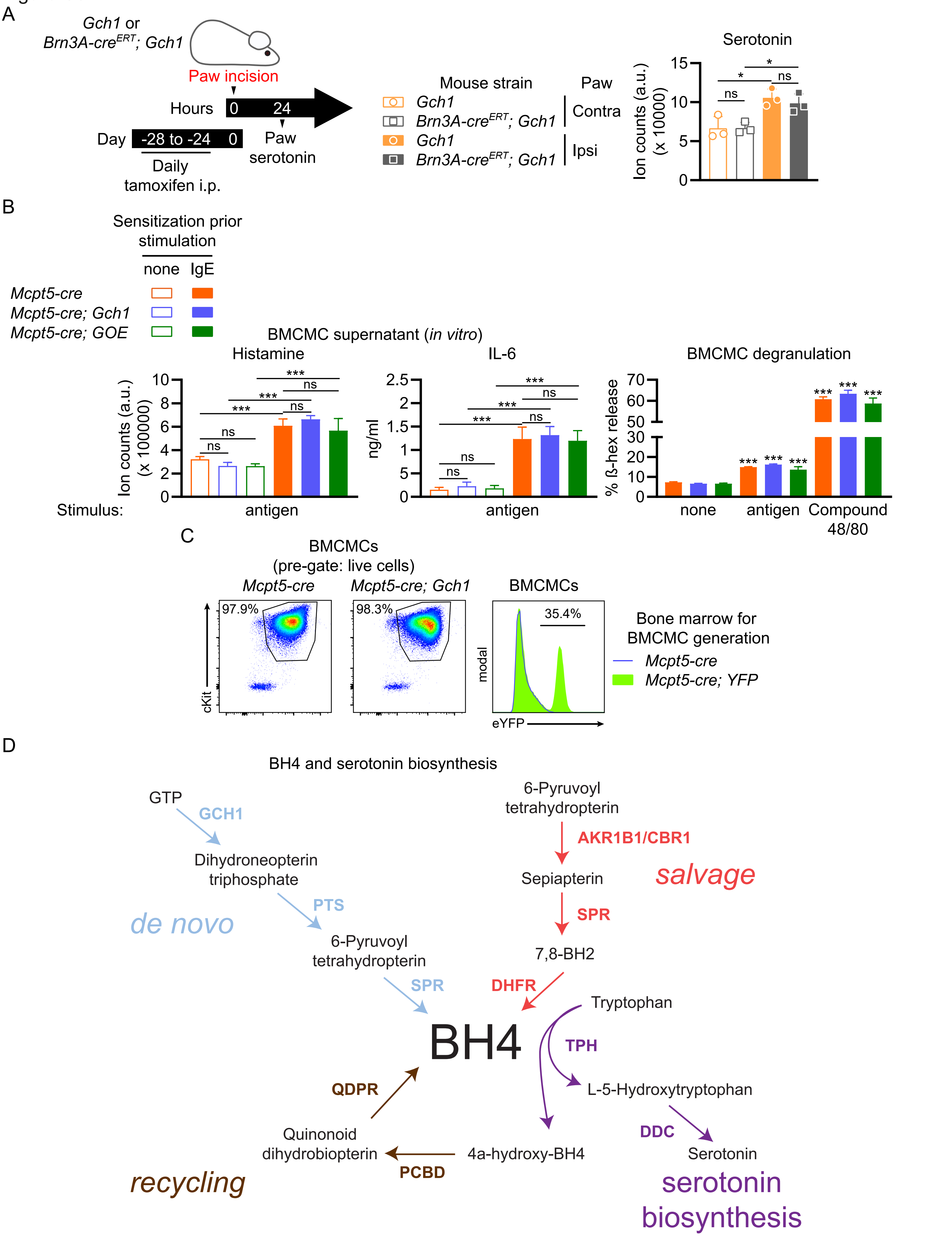
A
Gch1 or
Brn3A-cre^ERT; Gch1
Paw incision
Hours
0
24
Day
-28 to -24
0
Paw
serotonin
Daily
tamoxifen i.p.
Mouse strain
Paw
Gch1
Brn3A-cre^ERT; Gch1
Contra
Gch1
Brn3A-cre^ERT; Gch1
Ipsi
Serotonin
Ion counts (a.u.)
(x 10000)
ns
*
*
ns
B
Sensitization prior
stimulation
none
IgE
Mcpt5-cre
Mcpt5-cre; Gch1
Mcpt5-cre; GOE
BMCMC supernatant (in vitro)
Histamine
IL-6
BMCMC degranulation
Ion counts (a.u.)
(x 100000)
ng/ml
% ß-hex release
Stimulus:
antigen
antigen
none
antigen
Compound
48/80
C
BMCMCs
(pre-gate: live cells)
Mcpt5-cre
Mcpt5-cre; Gch1
BMCMCs
97.9%
98.3%
35.4%
cKit
modal
eYFP
Bone marrow for
BMCMC generation
Mcpt5-cre
Mcpt5-cre; YFP
D
BH4 and serotonin biosynthesis
GTP
GCH1
Dihydroneopterin
triphosphate
PTS
de novo
6-Pyruvoyl
tetrahydropterin
SPR
6-Pyruvoyl
tetrahydropterin
AKR1B1/CBR1
Sepiapterin
salvage
SPR
7,8-BH2
DHFR
Tryptophan
BH4
TPH
L-5-Hydroxytryptophan
QDPR
DDC
Quinonoid
dihydrobiopterin
4a-hydroxy-BH4
Serotonin
PCBD
recycling
serotonin
biosynthesis

### Figure S5. Paw serotonin levels of *Brn3A-cre$^{ERT}$; Gch1* mice, mediator release and degranulation of mouse bone marrow-derived cultured mast cells

(A) Left panel: experimental outline. *Gch1$^{flox/flox}$* (*Gch1*) mice or *Gch1* mice expressing tamoxifen-inducible cre recombinase in sensory neurons (*Brn3A-cre$^{ERT}$; Gch1$^{flox/flox}$*; *Brn3A-cre$^{ERT}$; Gch1*) mice were treated by daily intraperitoneal tamoxifen injections on 28 – 24 days before paw incision. Contra and ipsilateral (Ipsi) paw serotonin levels were assessed 24 hours after incision (n=3). (B) Bone marrow-derived cultured mast cells (BMCMCs) were generated from control (*Mcpt5-cre*) or *Mcpt5-cre; Gch1*, or *Mcpt5-cre; GOE* mice and activated by IgE and antigen, or Compound 48/80 for 1 hour, followed by analysis of histamine (by mass spectrometry), IL-6 (by ELISA) levels in the supernatant or relative ß-hexosaminidase (ß-hex) release (as degranulation surrogate). (C) Flow cytometry analysis of bone marrow-derived cultured mast cells (BMCMCs). Dot plots show mast cell gates (cKit$^{+}$/Fc$\varepsilon$RI$\alpha^{+}$) of cultures derived from *Mcpt5-cre* and *Mcpt5-cre; Gch1* (mast cell-specific *Gch1* deficient) mice. The histogram plot shows yellow fluorescent protein (eYFP) signal of BMCMC cultures derived from *Mcpt5-cre* and *Mcpt5-cre; YFP* reporter mice. (D) Schematic depicting the de novo, salvage and recycling arms of the BH4 pathway as well as the serotonin synthesis pathway. GTP, guanosine triphosphate; GCH1, GTP cyclohydrolase I; PTS, 6-Pyruvoyl tetrahydropterin synthase; SPR, sepiapterin reductase; AKR1, aldo-keto reductase family 1; CBR, carbonyl reductase family; DHFR, dihydrofolate reductase; QDPR, quinoid dihydropteridine reductase; PCDB, pterin-4alpha-carbinolamine dehydratase; TPH, tryptophan hydroxylase; DDC, dopa decarboxylase.

(A and B) One-Way ANOVA with Tukey's multiple comparisons test (asterisks above graphs in the "Degranulation" panel indicate statistical significance as compared to the respective unstimulated cells); * $P \leq 0.05$, *** $P \leq 0.001$, ns – not significant.

Figure S6

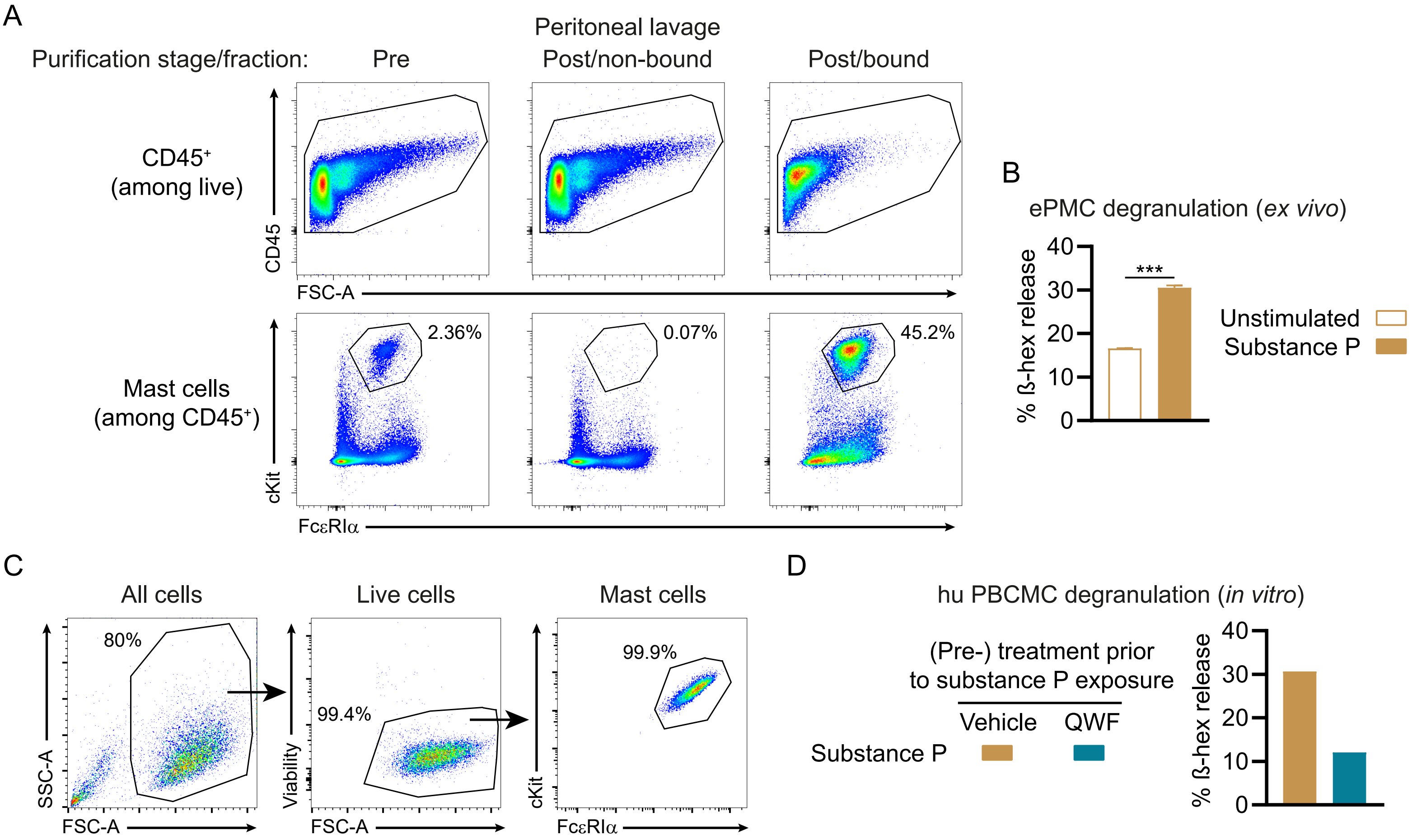
A
Peritoneal lavage
Purification stage/fraction:
Pre
Post/non-bound
Post/bound
CD45+ (among live)
CD45
FSC-A
Mast cells (among CD45+)
cKit
FcεRIα
2.36%
0.07%
45.2%
B
ePMC degranulation (ex vivo)
% ß-hex release
0
10
20
30
40
***
Unstimulated
Substance P
C
All cells
Live cells
Mast cells
80%
99.4%
99.9%
SSC-A
FSC-A
Viability
FSC-A
cKit
FcεRIα
D
hu PBCMC degranulation (in vitro)
(Pre-) treatment prior to substance P exposure
Vehicle
QWF
Substance P
% ß-hex release
0
10
20
30
40

#### Figure S6. Characterization and degranulation response of mouse peritoneal mast cells and human peripheral blood-derived cultured mast cells

(A) Flow cytometry analysis of peritoneal lavage fractions of C57BL/6 wild type mice of leukocytes ($CD45^+$) and mast cells ($cKit^+/Fc\varepsilon RI\alpha^+$) at different steps of magnetic beads-based mast cell enrichment: before enrichment (Pre); after enrichment (Post): non-bead-bound (non-bound) and bead-bound (bound); numbers indicate percentages of gated among all cells; (B) degranulation of enriched peritoneal mast cells (ePMCs) assessed by ß-hex release upon stimulation with 10 µM substance P for 1 hour. (C) Flow cytometry analysis of human peripheral blood-derived cultured mast cell (hu PBCMC) culture. (D) hu PBCMCs were pre-treated with vehicle or 100 µM QWF prior to simulation with (or without) 100 µM substance P for 1 hour. Degranulation was assessed by ß-hex release.

(B) Mann-Whitney test; n.d. – not detected (below detection limit); *** $P \leq 0.001$; error bars indicate SD.

### Supplemental table

#### Table S1. Cell populations and datasets used for *Gch1* expression analysis within the *ImmGen* platform

Table S1: *ImmGen/Gene Expression Commons* cell populations (related to Figure 1)

| **Cell type** | **Origin/subtype** | **GEXC model designation** | **Sorting Strategy** | **GEO dataset** | **GEO accession** | **Reference** |
|---|---|---|---|---|---|---|
| Dendritic cell | Lung[1] | DC_Lu_CD103$^+$ | CD11c$^+$/CD8a$^-$/CD11b$^{lo}$/ CD103$^+$ | GSE15907 | GSM538231; GSM538232; GSM538233; | (Gautier et al., 2012b; Miller et al., 2012) |
| Dendritic cell | Lung[2] | DC_Lu_CD11b$^+$ | CD11c$^+$/CD8a$^-$/CD11b$^{hi}$/ CD103$^-$/CD24$^+$ | GSE15907 | GSM854269; GSM854270; | (Gautier *et al.*, 2012b; Miller *et al.*, 2012) |
| Dendritic cell | Skin draining lymph node/CD4$^+$ | DC_SLN_4$^+$ | CD11c$^+$/CD8a$^-$/CD4$^+$/CD11b$^+$ | GSE15907 | GSM538245; GSM538246; GSM538247; | (Gautier *et al.*, 2012b; Miller *et al.*, 2012) |
| Dendritic cell | Skin draining lymph node/CD8$^+$ | DC_SLN_8$^+$ | CD11c$^+$/CD8a$^+$/CD4$^-$/CD11b$^-$ | GSE15907 | GSM538255; GSM538256; GSM538257; | (Gautier *et al.*, 2012b; Miller *et al.*, 2012) |
| Dendritic cell | Skin draining lymph node/CD11b$^+$ | DC_SLN8-4-11b+ | CD11c$^+$/CD8a$^-$/CD4$^-$/CD11b$^+$ | GSE15907 | GSM538271; GSM538272; GSM538273; | (Gautier *et al.*, 2012b; Miller *et al.*, 2012) |
| Dendritic cell | Skin draining lymph node/CD11b$^-$ | DC_SLN8-4-11b- | CD11c$^+$/CD8a$^-$/CD4$^-$/CD11b$^-$ | GSE15907 | GSM538268; GSM538269; GSM538270; | (Gautier *et al.*, 2012b; Miller *et al.*, 2012) |
| Dendritic cell | Epidermis/Langerhans cell | DC_SK Langer | CD45$^+$/MHCII$^+$/CD11c$^+$/CD11b$^+$ | GSE15907 | GSM538280; GSM538281; | (Gautier *et al.*, 2012b; Miller *et al.*, 2012) |

| | | | | | | |
|---|---|---|---|---|---|---|
| Macrophages | Peritoneum/F480$^{high}$ | Mϕ_Peritoneum F4/80$^{hi}$ | F480$^{hi}$/CD115$^{hi}$/CD11b$^{+}$/MHCII$^{-}$/CD11c$^{-}$ | GSE15907 | GSM605850; GSM605851; GSM605852; | (Gautier et al., 2012a; Heng and Painter, 2008) |
| Macrophages | Peritoneum/F480$^{low}$ | Mϕ_Peritoneum F4/80$^{lo}$ | B220$^{-}$/F480$^{lo}$/CD115$^{+}$/MHCII$^{+}$ | GSE15907 | GSM854294; GSM854295; GSM854296; | (Gautier *et al.*, 2012b) |
| Macrophages | Lung/interstitial | Mϕ_Lu inter. | MertK$^{+}$/CD64$^{+}$/CD11c$^{-}$/CD11b$^{+}$ | GSE37448 | GSM1136119; GSM1136120; GSM1136121; | (Heng and Painter, 2008) |
| Macrophages | Lung/alveolar[1] | Mϕ_Lu Alveol. | MertK$^{+}$/CD64$^{+}$/CD11c$^{+}$/CD11b$^{-}$ | GSE37448 | GSM1136122; GSM1136123; GSM1136124; | (Heng and Painter, 2008) |
| Macrophages | Lung/alveolar[2] | Mϕ_Lu_CD11b-CD11c+ | CD11c$^{hi}$/CD11b$^{-}$/CD103$^{-}$/MHCII$^{-}$/SiglecF$^{+}$. | GSE15907 | GSM538283; GSM538284; GSM538282; | (Gautier *et al.*, 2012b) |
| Macrophages | Adipose tissue | Mϕ_AT | CD45$^{+}$/CD64$^{+}$/Mertk$^{+}$ | GSE37448 | GSM1282109; GSM1282110; GSM1282111; | (Heng and Painter, 2008) |
| Macrophages | Skin draining lymph node | Mϕ_SLN Medull | CD169$^{+}$/CD11c$^{dim}$/CD11b$^{+}$/F480$^{+}$/B220$^{-}$/CD90$^{-}$/SiglecF$^{-}$/CD103$^{-}$/Ly6G$^{-}$ | GSE15907 | GSM854322; GSM854323; GSM854324; | (Heng and Painter, 2008) |
| Macrophages | Spleen red pulp | Mϕ_Sp red pulp | F480$^{hi}$/CD11b$^{lo}$/CD11c$^{-}$/autofluorescent | GSE15907 | GSM605853; GSM605854; GSM605855; | (Gautier *et al.*, 2012b) |
| Monocyte | Bone marrow | MO_BM 6C+II- | B220$^{-}$/CD3$^{-}$/CD115$^{+}$/Ly6C$^{+}$/MHCII$^{-}$ | GSE15907 | GSM854329; GSM854330; GSM854331; | (Gautier *et al.*, 2012b) |
| Monocyte | Blood[1] | MO_Blood 6+2- | B220$^{-}$/CD3$^{-}$/NK1.1$^{-}$/Ly6G$^{-}$/CD115$^{+}$/Ly6C$^{+}$/MHCII$^{-}$ | GSE37448 | GSM1136130; GSM1136131; | (Gautier *et al.*, 2012b) |

| | | | | | | |
|---|---|---|---|---|---|---|
| Neutrophil | Bone marrow | GN_Bone marrow | CD11b$^{+}$/ Ly6g$^{+}$ | GSE15907 | GSM605846; GSM605847; GSM605848; GSM605849; | (Ericson et al., 2014) |
| Neutrophil | Blood[1] | GN_Bl_Phase 1 | CD11b$^{+}$/Ly6g$^{+}$ | GSE15907 | GSM854306; GSM854307; GSM854308; | (Ericson *et al.*, 2014) |
| Neutrophil | Blood[2] | GN_Bl_Phase 2 | B220$^{-}$/CD4$^{-}$/SSC$^{hi}$/Ly6G$^{hi}$ | GSE37448 | GSM1282102; GSM1282103; GSM1282104; GSM1282105; | (Ericson *et al.*, 2014) |
| NK cell | Spleen | NK_Spleen_CD127- | CD45$^{+}$/NKp46$^{+}$/NK1.1$^{+}$/CD127$^{-}$/CD3$^{-}$/CD19$^{-}$/7-AAD$^{-}$ | GSE37448 | GSM1585330; GSM1585331; GSM1585332; | (Robinette et al., 2015) |
| NK cell | Liver | NK_Lv_CD49b+ | CD45$^{+}$/NKp46$^{+}$/NK1.1$^{+}$/CD49B$^{+}$/TRAIL$^{-}$/CD3$^{-}$/CD19$^{-}$/7-AAD$^{-}$ | GSE37448 | GSM1585333; GSM1585334; GSM1585335; | (Robinette *et al.*, 2015) |
| NK cell | Small intestine | NK_SI_CD127- | CD45$^{+}$/NKp46$^{+}$/RORgt$^{-}$/CD127$^{-}$/NK1.1$^{+}$/CD3$^{-}$/CD19$^{-}$/7-AAD$^{-}$ | GSE37448 | GSM1585336; GSM1585337; | (Robinette *et al.*, 2015) |
| ILC1 | Spleen | IL1_Sp_CD127+ | CD45$^{+}$/NKp46$^{+}$/NK1.1$^{+}$/CD27$^{+}$/CD127$^{+}$/CD3$^{-}$ CD19$^{-}$/7-AAD$^{-}$ | GSE37448 | GSM1585312; GSM1585313; GSM1585314; | (Robinette *et al.*, 2015) |
| ILC1 | Liver | IL1_Lv_CD49b- | CD45$^{+}$/NKp46$^{+}$/NK1.1$^{+}$/CD49B$^{-}$/TRAIL$^{+}$/CD3$^{-}$/CD19$^{-}$/7-AAD$^{-}$ | GSE37448 | GSM1585315; GSM1585316; GSM1585317; | (Robinette *et al.*, 2015) |
| ILC1 | Small intestine | IL1_SI_CD127+ | CD45$^{+}$/NKp46$^{+}$/RORgt-GFP$^{-}$/CD127$^{+}$/NK1.1$^{+}$/CD3$^{-}$/CD19$^{-}$/7-AAD$^{-}$ | GSE37448 | GSM1585318; GSM1585319; | (Robinette *et al.*, 2015) |
| ILC2 | Small intestine | IL2_SI_CD127+_Sca1+ | CD45$^{+}$/CD127$^{+}$/Sca1$^{+}$/ST2$^{+}$/KLRG1$^{+}$/CD3$^{-}$/CD19$^{-}$/7-AAD$^{-}$ | GSE37448 | GSM1585320; GSM1585321; | (Robinette *et al.*, 2015) |

| | | | | | | |
|---|---|---|---|---|---|---|
| ILC3 | Small intestine[1] | IL3_SI_NKp46+ | CD45$^+$/NKp46$^+$/RORgt-GFP$^{hi}$/CD3$^-$/CD19$^-$/7-AAD$^-$ | GSE37448 | GSM1585322; GSM1585323; GSM1585324; | (Robinette *et al.*, 2015) |
| ILC3 | Small intestine[2] | IL3_SI_NKp46-4- | CD45$^+$/RORgt-GFP$^+$/NKp46$^-$/CD4$^-$/CD3$^-$/CD19$^-$/7-AAD$^-$ | GSE37448 | GSM1585326; GSM1585327; GSM1585328; | (Robinette *et al.*, 2015) |
| ILC3 | Small intestine[3] | IL3_SI_NKp46-4+ | CD45$^+$/RORgt-GFP$^+$/NKp46$^-$/CD4$^+$/CD3$^-$/CD19$^-$/7-AAD$^-$ | GSE37448 | GSM1585325; GSM1585329; | (Robinette *et al.*, 2015) |
| Eosinophil | Adipose tissue | EO_AT | FSC$^{low}$/B220$^-$/CD11b$^+$/Siglec F$^+$ | GSE37448 | GSM1282093; | (Heng and Painter, 2008) |
| Eosinophil | Blood | EO_Bl | CD4$^-$/B220$^-$/SSC$^{hi}$/Siglec F$^+$ | GSE37448 | GSM1282094; GSM1282095; GSM1282096; | (Dwyer et al., 2016) |
| Mast cell | Trachea | MC_Tr | CD45$^+$/CD11b$^-$/CD11c$^-$/CD19$^-$/CD4$^-$/CD8$^-$/FcεR1α$^+$/CD117$^+$ | GSE37448 | GSM2112440; GSM2112441; GSM2112442; | (Dwyer *et al.*, 2016) |
| Mast cell | Tongue | MC_To | CD45$^+$/CD11b$^-$/CD11c$^-$/CD19$^-$/CD4$^-$/CD8$^-$/FcεR1α$^+$/CD117$^+$ | GSE37448 | GSM2112437; GSM2112438; GSM2112439; | (Dwyer *et al.*, 2016) |
| Mast cell | Skin | MC_Sk | CD45$^+$/CD11b$^-$/CD11c$^-$/CD19$^-$/CD4$^-$/CD8$^-$/FcεR1α$^+$/CD117$^+$ | GSE37448 | GSM2112434; GSM2112435; GSM2112436; | (Dwyer *et al.*, 2016) |
| Mast cell | Peritoneal | MC_PC | CD45$^+$/CD11b$^-$/CD11c$^-$/CD19$^-$/CD4$^-$/CD8$^-$/FcεR1α$^+$/CD117$^+$ | GSE37448 | GSM2112430; GSM2112428; GSM2112431; GSM2112432; GSM2112433; GSM2112429; | (Dwyer *et al.*, 2016) |
| Mast cell | Esophagus | MC_Es | CD45$^+$/CD11b$^-$/CD11c$^-$/CD19$^-$/CD4$^-$/CD8$^-$/FcεR1α$^+$/CD117$^+$ | GSE37448 | GSM2112426; GSM2112427; | (Dwyer *et al.*, 2016) |

| | | | | | | |
|---|---|---|---|---|---|---|
| Basophil | Spleen | BA_Sp | $CD3^-/CD19^-/NK1.1^-/CD117^-/Fc\varepsilon R1\alpha^+/CD49b^+$ | GSE37448 | GSM2112410; GSM2112411; GSM2112412; | (Dwyer *et al.*, 2016) |
| Basophil | Blood | BA_Bl | $CD4^-/CD8^-/CD19^-/NK1.1^-/CD49b^+/Fc\varepsilon R1\alpha^+$ | GSE37448 | GSM2112407; GSM2112408; GSM2112409; | (Dwyer *et al.*, 2016) |
